# Benchmarking antigen-aware inverse folding methods for antibody design

**DOI:** 10.1101/2025.08.05.668698

**Authors:** Bartosz Janusz, Dawid Chomicz, Sonia Wrobel, Pawel Dudzic, Adithya Polasa, Kyle Martin, Steven Darnell, Stephen R. Comeau, Konrad Krawczyk

## Abstract

Computational antibody design has seen many recent advances pioneered via the use of language models and advanced structure prediction tools. Developing a de novo antibody against a specific antigen requires structural awareness that most language models lack. A prominent class of machine learning methods combining the best of language model and structural worlds is inverse folding. This approach aims to predict a sequence that would fit a given structure. Such methods are now increasingly used to predict alternate sequences given a structure of a binder. It is known that, just like language models, such methods have certain predictive power in identifying binders. Here we performed a set of tests to reveal where, if at all, such methods provide value in the realistic setting of antibody discovery.

## Introduction

Antibodies play a critical role in the immune system and have become essential in therapeutics and diagnostics due to their highly specific antigen recognition capabilities. Engineering antibodies to enhance properties such as affinity, specificity, and stability has thus become an active research area, increasingly powered by machine learning and computational approaches (Bielska et al. 2025). Among these, inverse folding methods present a promising paradigm for antibody design (Li et al. 2024). Unlike traditional design approaches, which aim to determine a protein’s 3D structure based on a given sequence, inverse folding begins with a target structure and searches for sequences that best fold into that structure. This shift in focus has been particularly invigorated by advancements in structural prediction models like AlphaFold, which generates millions of predicted protein structures, fueling the training of inverse folding models (Varadi et al. 2022).

In the realm of antibody design, inverse folding methods offer distinct advantages by focusing on maintaining the native fold of an antibody structure, crucial for preserving its antigen-binding capabilities. Recent advancements in protein-generic inverse folding algorithms, such as ProteinMPNN (Dauparas et al. 2022) and ESM-IF (Shanker et al. 2024), have achieved high sequence recovery and structural fidelity, establishing a foundation for antibody-specific inverse folding models. For instance, AbMPNN (Dreyer et al. 2023) and AntiFold (Høie et al. 2025) have fine-tuned these foundational methods using antibody-focused datasets, yielding notable improvements in sequence recovery, structural accuracy, and binding affinity predictions across antibody complementarity-determining regions (CDRs).

Despite these innovations, evaluating the effectiveness of inverse folding methods for antibody design poses unique challenges. Current evaluation metrics for generative models often rely on in silico criteria, which are split into sequence-based metrics, such as amino acid recovery (AAR), and structure-based metrics, such as root-mean-square deviation (RMSD) from target structures. Previous study into benchmarking antibody inverse folding methods chiefly focused on such in-silico metrics (Li et al. 2024). Using public datasets, this study revealed important shortcomings of these methods, such as weak correlations with affinity and limited usage of antigen information, in contradiction to previous findings (Shanker et al. 2024). Such metrics, though useful, might not be directly transferable to antibody discovery campaigns functional performance. Furthermore, structure-based confidence scores like predicted Local Distance Difference Test (pLDDT) and interface-predicted Template Modeling (ipTM), while valuable for structural validation, are inadequate proxies for binding efficacy in antibodies. Physics-based metrics, though theoretically promising due to their consideration of biophysical properties, face high computational demands and have shown weak correlations with experimental affinity measurements (Hummer et al. 2023).

In this work, we address these limitations by benchmarking state-of-the-art inverse folding models specifically for antibody design. We perform evaluation on three diverse experimental datasets, focusing on realistic design applications, including a proper hold out in the form of our structure-inform AbDesign dataset (Janusz et al. 2025). This benchmark confirms that ML methods such as inverse folding can be used to enrich binders from a large pool of potential ones. Nonetheless, our results show that the methods make very limited use of the antigen, which leaves a clear direction for improvement.

## Methods

### AbDesign DB dataset

The AbDesign DB dataset was generated by selecting seven antigens, with each one being recognized by two distinct antibodies. Some of these targets were therapeutically significant, like VEGF and PD-1, while others were common antigens that are frequently crystallized in the PDB. The CDR-H3 regions had a median length of 12 residues. However, our primary focus was not specifically on therapeutic antibodies or relevant targets.

We identified interactions between the CDR-H3 regions and antigen residues, defining these contacts as cases where heavy atoms were within a distance of 4.5Å. These contact positions were subsequently chosen for point mutations. The aim was to maximize the number of single mutants and distinct mutation sites while keeping the total number of variants capped at 96 antibodies per plate per target. To achieve this, mutations were distributed automatically to ensure a wide range of positions were covered, avoiding a bias towards any specific site. Each mutation set was then manually reviewed to exclude non-informative or irrelevant mutations, such as the introduction of cysteines. This resulted in a total of 658 mutations, averaging 47 mutations per antibody.

We expect this dataset to be particularly useful in affinity maturation experiments, where mutations need to be introduced without affecting wild-type binding. To support this, we modeled each antibody structure using ABodyBuilder2 (Abanades et al. 2023) and optimized the structures with OpenMM. Each file was labeled with its specific mutation. The structural data can be cross-referenced with sequence information and ELISA ratios, making it suitable for enhanced machine learning analyses.

### Anti-PD1 dataset

The original dataset was composed of 59 unique heavy and light sequences of binding antibodies only. Additional 59 non-binder sequences were generated in the following manner: for each heavy binder sequence, replace part of CDR3 (leave 2 AAs at the beginning, 3 at the end) with a random AA sequence (excluding cysteine) matching the original length. The sequences have varying lengths (Supplementary Figure 1).

### Trastuzumab HER2 Large

The HER2-aff-large dataset consists of over half a million Trastuzumab variants categorized into three affinity classes—’high,’ ‘medium,’ and ‘low’—based on their binding strength to HER2 (Chinery et al. 2024). These variants were derived by limiting mutations to the IMGT positions between 107 and 116 of the antibody’s heavy chain. The dataset is an extension of previous DMS (Deep Mutational Scanning) results from Mason et al. and includes 178,160 high-affinity, 196,392 medium-affinity, and 171,732 low-affinity binders.

A small percentage of overlap between affinity classes was observed, with 1.1% of sequences classified as both ‘high’ and ‘medium’ binders, and 2.9% found in both ‘high’ and ‘low’ categories. By addressing these overlaps, the dataset was refined to 530,357 sequences, resulting in a 33.6% class imbalance. Further refining by removing all overlapping sequences entirely reduced the dataset to 524,346 sequences with a class imbalance of 32.8%. CDRH3 residue distributions can be seen in Supplementary Figures 2 and 3.

The dataset was used to analyze binding behaviors, with clustering analysis showing that the ‘medium’ and ‘low’ classes grouped with negative binders from the original Mason et al. study. Classification models predicted high binding probabilities for the ‘high’ affinity class and low probabilities for the ‘medium’ and ‘low’ affinity groups. For binary classification, the high-affinity sequences were labeled as positive binders, and the medium- and low-affinity sequences were grouped as negative binders, aligning with the goal of selecting high-affinity antibodies during the optimization process.

### Inverse Folding methods benchmarked

Four inverse-folding models were selected in this study, two protein generic ones (ESM-IF and ProteinMPNN) and their two antibody-focused versions (AntiFold and AbMPNN).

The ESM-IF1 (Hsu et al. 2022) inverse folding model is a computational tool developed by Meta AI as part of the Evolutionary Scale Modeling (ESM) series, which focuses on protein structure prediction and analysis. ESM-IF1 is specifically designed for protein inverse folding, a task that involves predicting a protein sequence given its three-dimensional (3D) structure, rather than the more common task of predicting the structure from a sequence. It is based on a GVP network trained on millions of structures from the AlphaFold2 Database.

AntiFold (Høie et al. 2025) is fine-tuned from the ESM-IF1 model with improved sequence recovery in complementarity-determining regions (CDRs), which are crucial for antigen recognition. It significantly improves upon existing models in this area, achieving a mean root-mean-square deviation (RMSD) of 0.95 for CDR regions, compared to higher RMSD values from other models like AbMPNN and ProteinMPNN.

ProteinMPNN (Dauparas et al. 2022) is a deep learning-based method for protein sequence design in the Baker Lab. It is based on a message passing neural network, and it was trained on the crystal structures from the PDB rather than model structures, as was the case with ESM-IF.

The AbMPNN (Dreyer et al. 2023) model is a fine-tuned, antibody-specific graph neural network model based on ProteinMPNN, designed to predict and design antibody sequences. It focuses on the variable CDR loops, particularly CDR-H3, which is vital for antigen recognition. The model is trained on large-scale antibody datasets such as SAbDab and OAS, as well as predicted structures from ABodyBuilder2, to provide state-of-the-art performance in antibody sequence recovery and design.

### Perplexity definition

In all benchmarked models, perplexity is defined as the exponential of cross entropy loss, averaged over all (or subset) residues in a scored sequence.

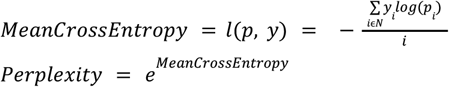

where *N* is sequence length.

## Results

### Overview of experiments

We tested four inverse folding models: ProteinMPNN, AbMPNN, ESM-IF, and AntiFold. ProteinMPNN and ESM-IF are state-of-the-art protein inverse folding models, whereas AbMPNN and ESM-IF are their antibody versions, respectively. The choice of the datasets reflected state-of-the-art & allowed us to test whether fine-tuning on antibodies brings any benefits in actual antibody discovery tasks beyond amino acid recovery, which is the objective task the models are trained on.

We performed tests on three datasets, HER2, BI-PD1, and NaturalAntibody AbDesign DB. The HER2 dataset is a public resource with 500,000 trastuzumab variants. We employed this dataset as a sanity check to see whether in our hands, we could get the inverse folding models to demonstrate their basic reported function - correlate with higher affinity binders. The BI-PD1 dataset was internally provided by Boehringer-Ingelheim (BI) with PD1 binders and non-binders. The experiment on HER2, which was supposed to separate higher from lower affinity binders, was supposed to be repeated on a real dataset used for antibody discovery. The final dataset, NaturalAntibody AbDesign DB, is a heterogeneous dataset of CDR-H3 mutants of existing structures. This was the final experimental test on how the methods fare in seemingly easy tasks of suggesting binding CDR-H3 mutations.

### Benchmarking enrichment of Trastuzumab variants against HER2

We benchmarked four inverse folding models—ESM-IF, Antifold (ESM-IF fine-tuned on antibodies), ProteinMPNN, and AbMPNN (ProteinMPNN fine-tuned on antibodies)—using the HER2-aff-large dataset containing over 500,000 CDRH3 sequences, labeled with three affinity categories: high, medium, and low. Our goal was to evaluate each model’s ability to accurately discriminate between high-affinity binders, medium-affinity binders, and low-affinity non-binders. This was designed to simulate a realistic antibody design scenario when we are faced with a set of sequences with unknown binding labels. We test whether a model, in this case inverse folding, has capacity to enrich top binders. It must be noted that in no case here do we talk about zero-shot predictions, as HER2 and Trastuzumab were likely seen by all models involved.

First, we performed a simple check to see whether sorting the models has any predictive power in separating binders from nonbinders on this dataset. We plotted histograms of the perplexity scores across these affinity groups under two structural conditioning scenarios: native structures and structures modeled with ABodyBuilder2 (Figure 1). It is clear that regardless of the model or whether we employ the native structure, there is a certain degree of separation between high/medium/low affinity binders. The main question is whether this slight degree of separation translates to an enrichment that is useful in an antibody engineering scenario.

**Figure 1.**
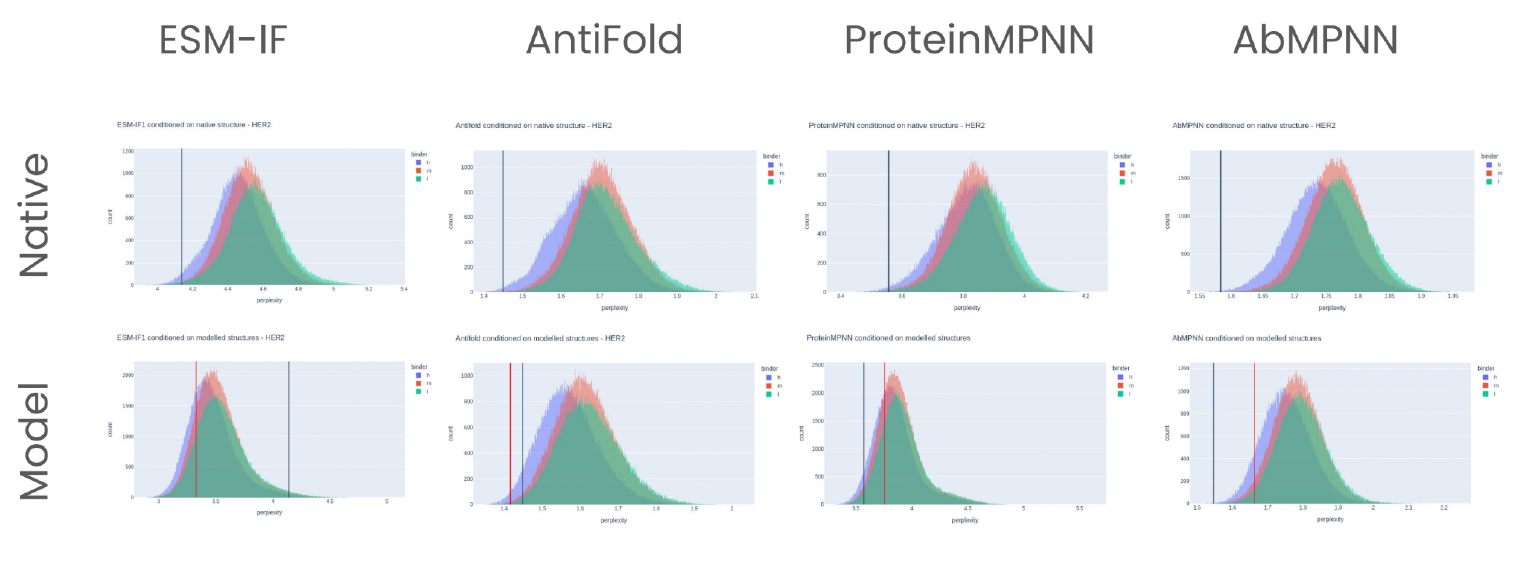
Sorting sequences by perplexity provides a degree of separation of binders and nonbinders. For each sequence in the HER-2 datasets, we calculated perplexity using one of four models ESM-IF, ProteinMPNN, AntiFold, or AbMPNN conditioned either on the native Trastuzumab structure or on the ABodyBuilder2 model. Black lines indicate native sequence (Trastuzumab) conditioned on native structure.

Overall, all four models demonstrated varying levels of capacity to distinguish between binders and non-binders based on perplexity distributions. Specifically, AbMPNN consistently emerged as the top performer, offering superior separation between high and low binders. Antifold also displayed strong separation capability, outperforming ESM-IF and ProteinMPNN, particularly in clearly distinguishing high-affinity sequences. In cases of ProteinMPNN and AntiFold, the top plate would have between 70-80% binders (Figures 2, 3).

**Figure 2.**
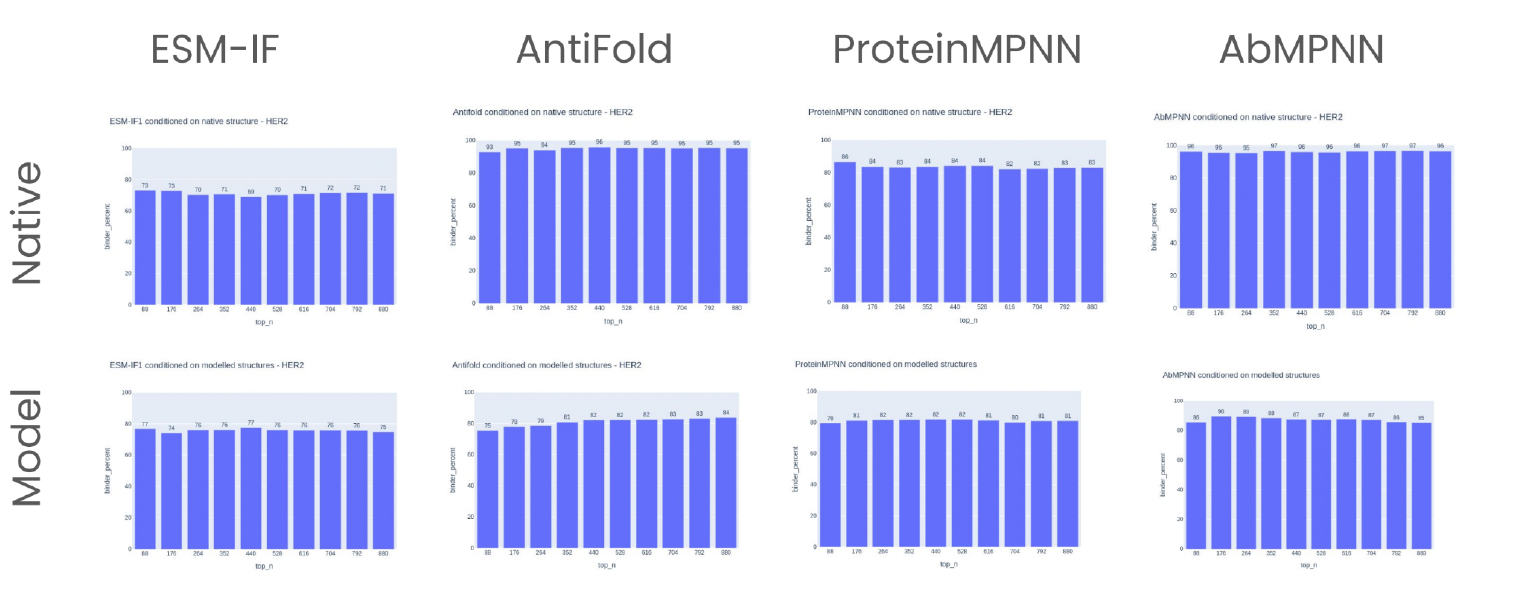
Sorting the top-N sequences based on perplexity - medium are treated as binders. The bins are multiples of 88, simulating a scenario where we get to select top N sequences from a larger set of unknown binders/nonbinders and can test them in batches of 88.

**Figure 3.**
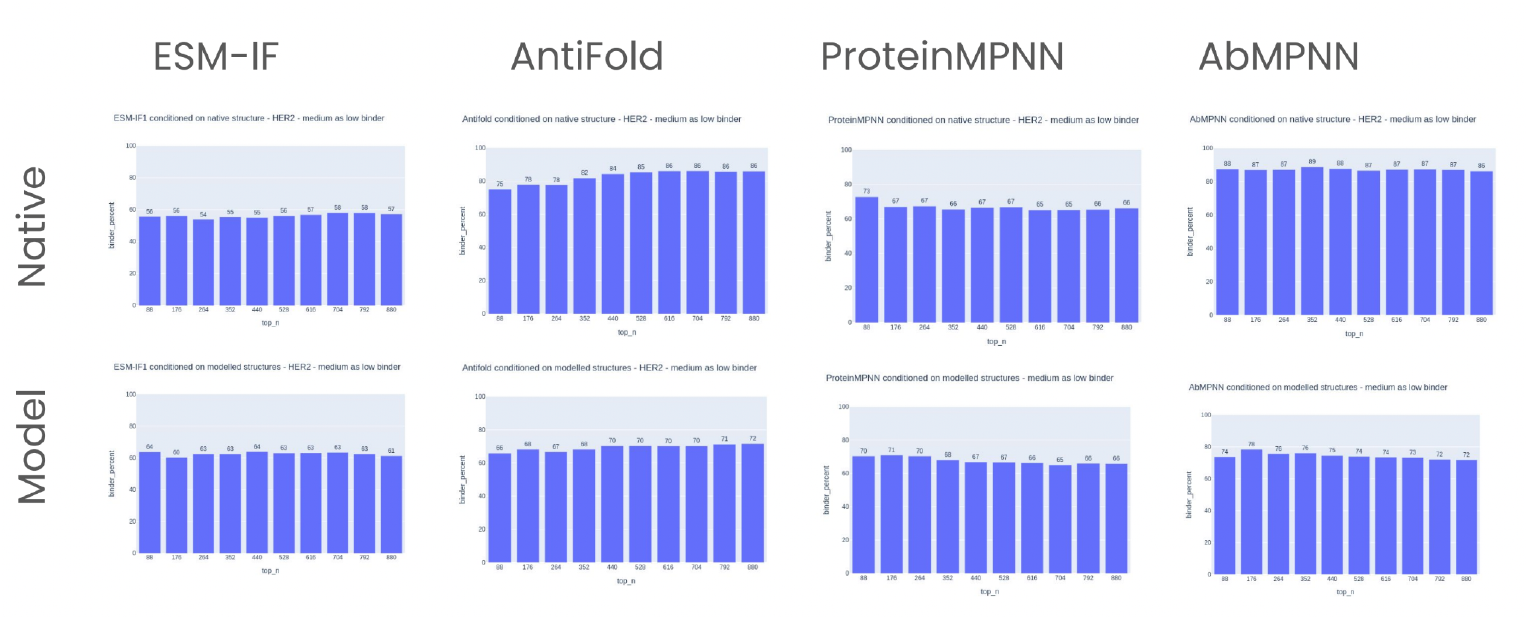
Sorting the top-N sequences based on perplexity - medium are treated as nonbinders. To address the antibody engineering question, we calculated the binding/nonbinding labels of the top-N sequences as sorted by perplexity (Figures 2, 3). The analysis was based on microplate granularity, assuming a standard one with 88 wells. The question we answered was: how many binders can we expect in 1 plate, if we choose to test sequences with the lowest perplexity first? Hence, we present charts with a high binder percentage among top N sequences, N being a multiple of 88. We tested that hypothesis on two variants of HER2-aff-large dataset 1. keep high and low binders (Figure 2), and 2. remove medium affinity binders, treat medium affinity binders as low (Figure 3).

Native structure conditioning unsurprisingly resulted in more precise distinctions between binding affinity groups across all models. When conditioning was based on ABodyBuilder2-modeled structures, we observed a general decline between 5-10 percentage points in the accuracy of binder/non-binder separation (Figure 2, 3). The fine-tuned antibody-specific models (Antifold and AbMPNN) were especially sensitive to structural conditioning, showing a notable performance drop when native structures were replaced by modeled structures. Despite this reduction, AbMPNN maintained the highest overall accuracy even when conditioned on modeled structures.

Considering binder enrichment among top-ranked sequences (based on lowest perplexity), AbMPNN consistently demonstrated the highest performance, achieving an enrichment rate of 96% when distinguishing between high and low binders, which remained robust at 88% even when medium-affinity sequences were grouped with non-binders. AntiFold closely followed, achieving 93% enrichment for high vs. low binders, decreasing slightly when medium binders were treated as non-binders (75% for the top sequences). ProteinMPNN outperformed ESM-IF but fell short of the antibody-specific models, maintaining a competitive enrichment rate of 86% (high vs. low binders) and 73% when medium binders were considered non-binders.

Treating medium binders as non-binders significantly impacted all models’ predictive accuracy, reducing their binder enrichment levels (Figure 3). The greatest resilience to this classification change was observed in AbMPNN, followed by Antifold, while ProteinMPNN and ESM-IF showed more noticeable performance declines. This shows that simplifying a continuous problem of affinity prediction to binary labels and categories is very problematic for models.

In summary, AbMPNN emerged as the most effective model for distinguishing high-affinity binders from non-binders within the HER2 dataset. This example shows the benefits of domain-specific modeling as both models did better than their base models, ESM-IF and ProteinMPNN. A big caveat though, is that the Trastuzumab crystal structure has likely been used in training, as indicated by the drop of modeling accuracy. Nonetheless, even for well-characterized targets such as HER-2, the protocol of employing inverse folding to sort through NGS or sequence-generated results appears to have a degree of benefit in providing enrichment of binders in the top plates.

### Anti-PD1 - internal BI dataset

Following our assessment of inverse folding methods on the publicly available HER2 dataset, we investigated the applicability of these approaches in an industrial context using an internal BI dataset comprising PD1 binder and non-binder antibody pairs. This dataset includes 59 distinct heavy and light chain sequences of varying lengths. Due to the absence of native structural information, we relied entirely on structures modeled using ABodyBuilder2 for all analyses. Evaluation was conducted by conditioning heavy chain sequences on their corresponding modeled structures, with a balanced dataset (50% binders, 50% non-binders).

Initially, we examined the perplexity score distributions of binders and non-binders across all four models: ESM-IF, Antifold, ProteinMPNN, and AbMPNN (Figure 4). Consistent with our HER2 observations, there was modest enrichment of binders at the lower perplexity scores, although the separation was less pronounced due to the smaller dataset size.

**Figure 4.**
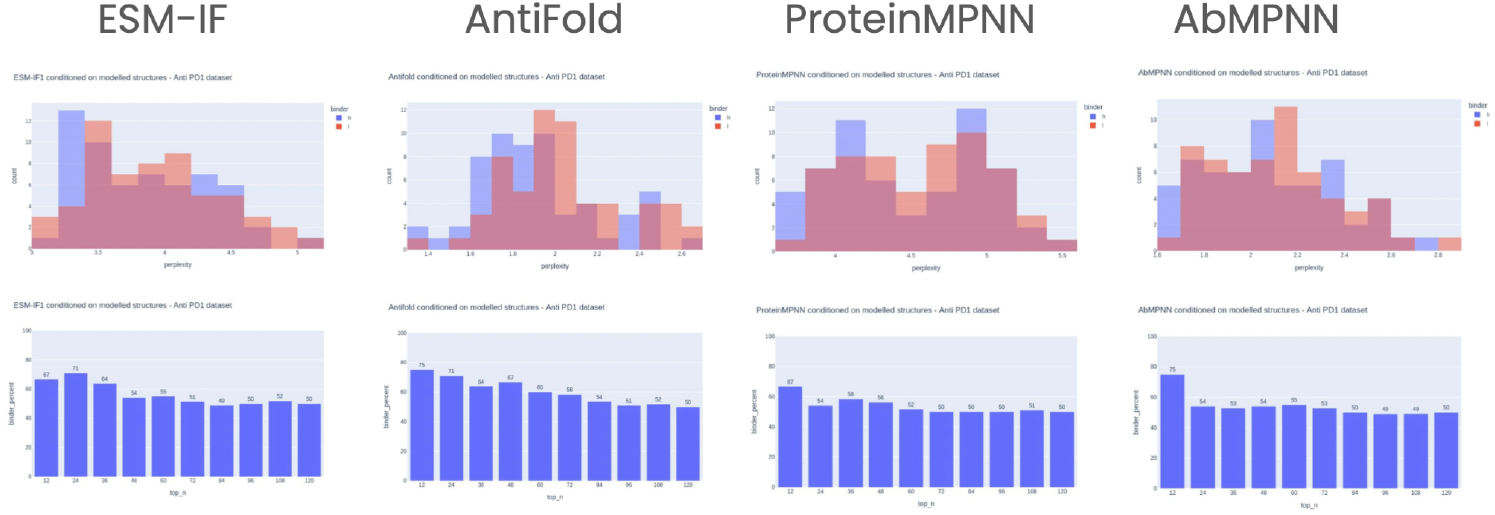
Separation of binders vs non-binders for the BI-PD1 dataset. **Top:** binders (h) and non-binders (l) histograms by their perplexities. Bottom: binders in top N sequences scored by perplexity, taken in batches of 12.

Among the tested models, Antifold provided the clearest differentiation between binder and non-binder sequences based on perplexity distributions, achieving a stable binder enrichment of 75% among the top 12 sequences, steadily declining toward the expected baseline as more sequences were considered. This indicates that Antifold reliably captures sequence features associated with higher affinity, even with limited data.

AbMPNN initially matched Antifold’s performance with 75% enrichment among the top 12 sequences. However, enrichment declined rapidly in larger sequence groups, suggesting lower stability and less robust separation between binder types. The stochastic nature inherent to ProteinMPNN-based models likely contributed to this performance variability, highlighting potential limitations when applied to smaller datasets.

ESM-IF, despite being a general-purpose model, surprisingly showed competitive initial performance, reaching 71% enrichment in the top 24 sequences. Nonetheless, this advantage quickly diminished beyond this threshold, dropping to about 54% at larger selections and demonstrating relatively limited ability to maintain robust binder enrichment across a wider dataset.

ProteinMPNN exhibited the weakest performance overall, with no clear separation between binders and non-binders. The maximum observed enrichment was only 67%, and enrichment was inconsistent, with binder peaks appearing at both moderate and higher perplexity scores. This further underscores the advantage of antibody-specific models, particularly in datasets with limited sequence diversity and size.

Collectively, our analysis demonstrates that antibody-specific inverse folding models, particularly Antifold, perform more reliably and effectively in discriminating binders from non-binders in industrial-scale datasets. General protein models and stochastic antibody-specific models (e.g., AbMPNN) exhibit variable performance, underscoring the need for further optimization or larger datasets for consistent predictive accuracy.

### Performance on AbDesign DB

The previous HER2 and PD1 datasets served to demonstrate that there is some predictive power in applying inverse folding models to surface antibodies with a higher likelihood to bind, out of a set of known binders/non-binders. In these datasets, however, it was impossible to gauge to what extent the antigen affects the performance of the models. In the case of HER2, the mutants were typically multiple mutations away from the parent Trastuzumab, so modeling the complex between it and HER2 would be unreliable in the first place. In the case of PD1, we had no way of modeling the interaction between the sequences and PD1 as antibody-antigen docking is still an open problem (Hitawala and Gray 2024).

To address this, we employed the NaturalAntibody AbDesign DB. The resource consists of 14 antibody-antigen complexes, seven targets, and two antibodies per target. There are ca. 700 point mutants of CDR-H3, measured by ELISA. Therefore, we know the precise structural configuration, the mutations, and the resulting effect on binding. The effect of mutation is measured as a ratio of ELISA binding of the mutant to the wild type. Therefore, values close to 1.0 indicate maintenance of binding at least as good as the parent, whereas values closer to 0.0 indicate loss in binding.

We applied the inverse folding models to it to check whether the single-point mutation effect can be predicted. We performed conditioning on the model of the antibody as well as native structure to see the effects. For antigen-specific conditioning, we only employed ESM-IF and its antibody-specific version, AntiFold as ProteinMPNN was unable to handle the antigen information.

For each of 14 complexes, we plotted the perplexity score versus the ELISA ratio (Supplementary Figure 4-11). An ideal correlation would be an increasing linear slope from left to right, however, in many cases, the tendency was negative. We collated the overall correlations in Table 1. These indicate that there appears to be a major improvement whether a native structure is used over a model. Furthermore, the antibody-specific version AntiFold, performs better than ESM-IF, retaining some predictive power in the models, though a very weak one. There appears to be a minor improvement in correlation, while using antigen, but it is very weak, if any. We recognize that our proxy binding metric of ELISA ratio might be indicative in a more binary fashion whether there is maintenance of binding. Otherwise, the correlation, as would be expected with affinity, is quite tenuous.

**Table 1.** Comparison of correlations between perplexities and the ELISA ratio on NaturalAntibody AbDesign dataset.

| model | conditioning type | multi chain pearson | single chain pearson |
| --- | --- | --- | --- |
| Antifold | modeled | 0.086105 | 0.119650 |
|  | native | 0.239855 | 0.247221 |
| ESM-IF | modeled | -0.012940 | 0.036561 |
|  | native | 0.181204 | 0.189677 |

To account for unreliability of the correlation scheme, we reformulated the task in a binary fashion, defining binding as everything above 1.0 and non-binding as below 1.0 (Supplementary Figures 12-19). This was supposed to simulate a scenario if we enumerated the mutations in question, calculated perplexities and only kept as binding those with scores above the wild type 1.0 If the scores have any predictive power, the box plot for each of the PDBs should be tending towards zero towards the left, and towards one to the right. We counted how many such PDBs had tendency in the direction of making correct predictions and called it a success, with the results in Table 2. Again, the performance of native structures greatly outperforms the models, underpinning the need for better structural predictions. Including the antigen in the equation did not appear to improve matters.

**Table 2.** Success rate of picking the mutants that maintain binding.

| model | conditioning type | multi chain success (n=14) | single chain success (n=14) |
| --- | --- | --- | --- |
| Antifold | modeled | 5 | 5 |
|  | native | 10 | 10 |
| ESM-IF | modeled | 3 | 4 |
|  | native | 11 | 9 |

In conclusion, the inverse folding scheme can be used, when starting from a native binder structure. If a model is used the results will not be as reliable. There appears to be a very weak effect of antigen on the predictions which leaves a lot of room for improvement of the methods.

### Testing the efficacy of sampling and oracle schemes

The previous analyses all focused on pre-existing datasets to check how the machine learning scores would sort them. Though this shows whether a model has a positive tendency to enrich for favorable sequences, it does not tell us whether such better sequences would actually be generated by the model.

We checked if sampling using inverse-folding models produces binder sequences. We trained oracle classifiers (CNN architecture as previously from a DMS study (Mason et al. 2019)) on the HER2 Large dataset in three configurations: only on CDRH3 sequences treating medium binders as low (Supplementary Figures 20 and 21) only on CDRH3 removing medium binders (Supplementary Figure 22) and finally on full heavy sequences using medium as low binders (Supplementary Figure 23). Then it was used to determine if the sampled sequence is a binder or a non-binder. We also prepared a dataset of 10,000 random CDRH3 for a sanity check of classifiers. Only the CDRH3 sequence was used for sampling and classification.

We noticed that it is problematic to sample sequences at low temperatures - so close to the original binder sequence. The sampling happens by sampling the multivariate distribution for each residue produced by the model. Low temperatures have naturally biased distribution, which is the reason the multivariate oscillates around the same answers. The models produce low amounts of unique sequences (around 0.1% and less on temp 1 and fine tuned models). Though we note that MPNN-based models give around 10% more binders and produce more diverse repertoires quicker: over 100 unique sequences vs 36 by Antifold in similar time.

A practical solution we found to this can be mitigated by different sampling strategies, such as enumeration and sorting by perplexity. Here, the ‘temperature’ could be implicitly incorporated by further filtering of the sequences to only the ones that are a certain, lower, sequence identity away from the parental sequence.

The other issue we found was to do with the oracles themselves. Training a model on a large number of binders-non binders has been independently shown to produce models that can be called binder-non-binders experimentally. However, in our experiments, we controlled with ‘random’ sequences as the negative, non-binding class. Models show great variation in prediction accuracy depending on whether the medium-affinity binders are included as positive class (Supplementary Figures 20-27).

The most worrisome though, were the sanity checks, where we checked how many randomly generated CDRH3s would be classified as binders. In all cases, a large proportion is classified as such - 5.6%, 38.0%, 10.6%. However, we would treat the medium binders, it is quite extraordinary that as many as 5.6% random CDR-H3 sequences would still maintain binding. Though that was not verified experimentally. Please note that the pie charts would be identical between ESM-IF/Antifold and ProteinMPNN/AbMPNN because the sequences were generated randomly, and posteriori we checked how likely the model would have generated them.

### Sampled datasets statistics

The goal of the sampling procedure was to have 1000 unique sequences for each model-temperature combination. However, because of extremely long sampling time at lower temperatures (models generating a large number of non-unique sequences), we stopped the sampling with much smaller numbers of sequences (Table 3). We calculated sequence identity between each sampled CDRH3 sequence and the native one taken from Trastuzumab. We noticed that identity drops with rising temperature, which is the expected behavior, but to our surprise, not to a big extent (Figure 5). Residue distributions confirmed that even with drastic temperature differences (0.1 vs 1.0), residues generated on some positions are highly conserved (Figure 6).

**Table 3:** Sampled datasets counts. Counts of unique sequence sets sampled using 4 models on different temperature settings. The higher the temperature, the more diverse sequences a model produces, hence easier (faster) to generate unique examples. For the lowest (0.1) temperature, Antifold was generating identical sequences most of the time, thus such low cardinality (36) of the corresponding dataset.

|  | Sampling temperature |  |  |  |  |  |
| --- | --- | --- | --- | --- | --- | --- |
| Model name | 0.1 | 0.2 | 0.4 | 0.6 | 0.8 | 1 |
| AbMPNN | 114 | 553 | 774 | 841 | 979 | 1000 |
| Antifold | 36 | 200 | 500 | 1000 | 964 | 999 |
| ESM-IF | 336 | 500 | 1000 | 999 | 1000 | 1000 |
| ProteinMPNN | 194 | 470 | 944 | 996 | 1000 | 1000 |

**Figure 5:**
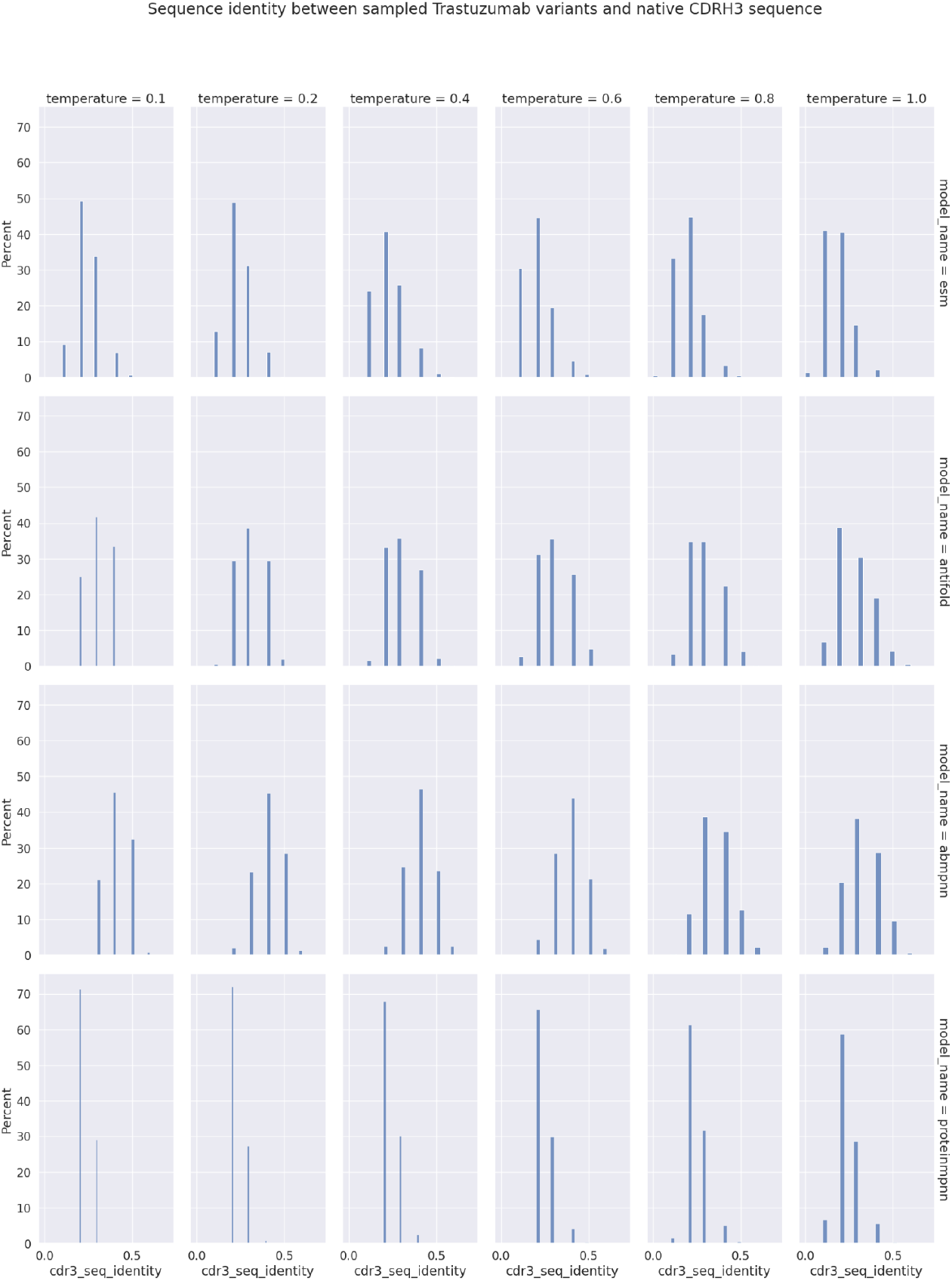
Sequence identity distributions between sampled and native CDRH3 sequences. Each model tends to output similarly distant sequences across different temperature settings. Sequence identity drops at most ~10% between 0.1 and 1.0 temperature for ESM-IF and ProteinMPNN. For antibody-specific models the drop is even less (~5%).

**Figure 6:**
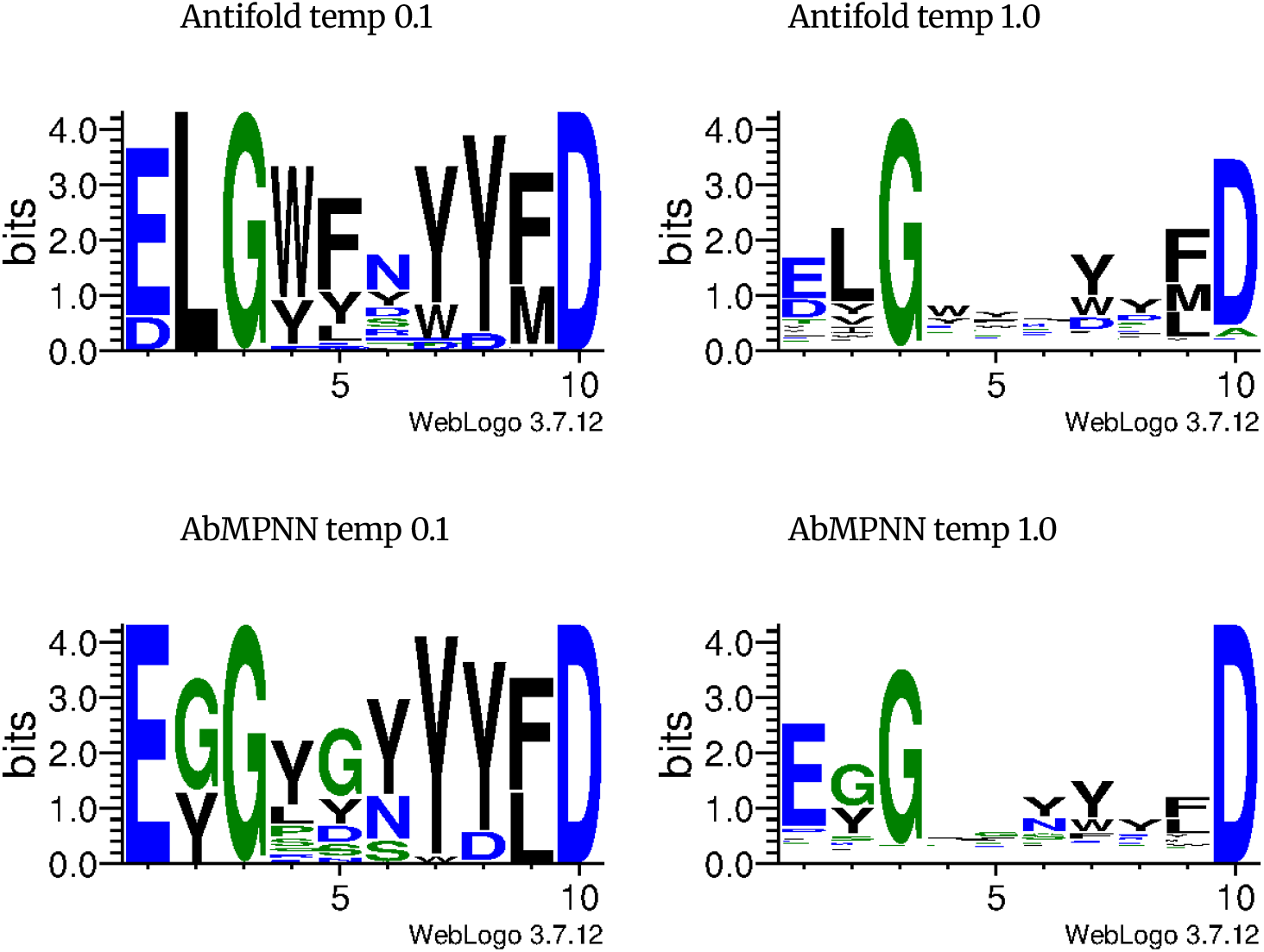
WebLogo residue distributions for Antifold and AbMPNN sampled CDRH3 sequences. We can see that 3rd (G) and 10th (D) positions are highly conserved for both AntiFold and AbMPNN.

## Discussion

Generative methods hold great promise in accelerating the costly and arduous experimental design process of biologics discovery. To this end, we benchmarked structure-aware generative methods of the inverse-folding disposition. This was dictated by desire to see whether these methods are useful in a realistic discovery setting and to what extent the antigen influences the predictions.

Our analysis confirms observations of other studies that, given a set of antibodies, these methods are useful in enriching for the higher-affinity antibodies (Hie et al. 2024). They do so in a manner agnostic to the antigen, and they can operate on a model of the antibody sequence. Therefore, they are suitable for NGS datasets from display or immunizations, as typically used in the course of discovery campaigns (Erasmus et al. 2023).

We tested the methods on our own internal dataset that was supposed to reveal to what extent the antigen information is taken into account. This test revealed that the antigen does not inform the predictions a great deal, if at all, as confirmed by other studies on this topic (Uçar, Malherbe, and Gonzalez 2024).

Altogether, our findings suggest that the inverse folding models are moderately useful in an industrial setting for filtering NGS datasets. There is however large room for improvement to bias the datasets in the direction of taking the binding interface into account.

