## Supplementary Information for "Benchmarking antigen-aware inverse folding methods for antibody design"

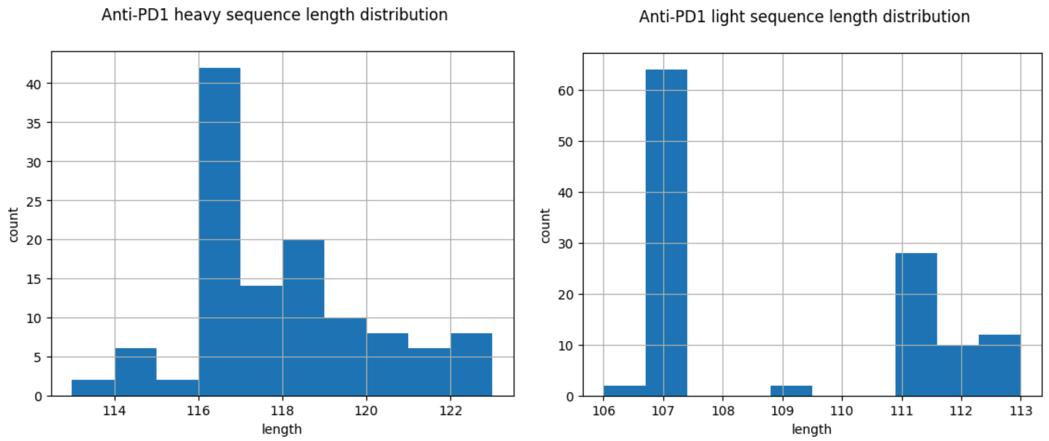


**Supplementary Figure 1: Anti-PD1 heavy (Left) and light (Right) variable region sequence length distribution.**

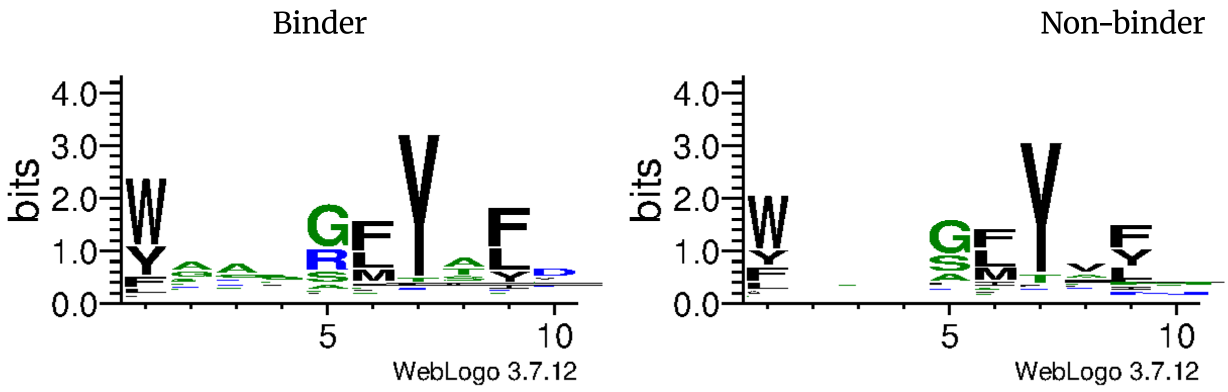


**Supplementary Figure 2: WebLogo residue distributions for HER2-large-aff CDRH3 sequences -** medium labeled as low binders, binder (Left) and non-binder (Right). We can see 7th position is highly conserved.

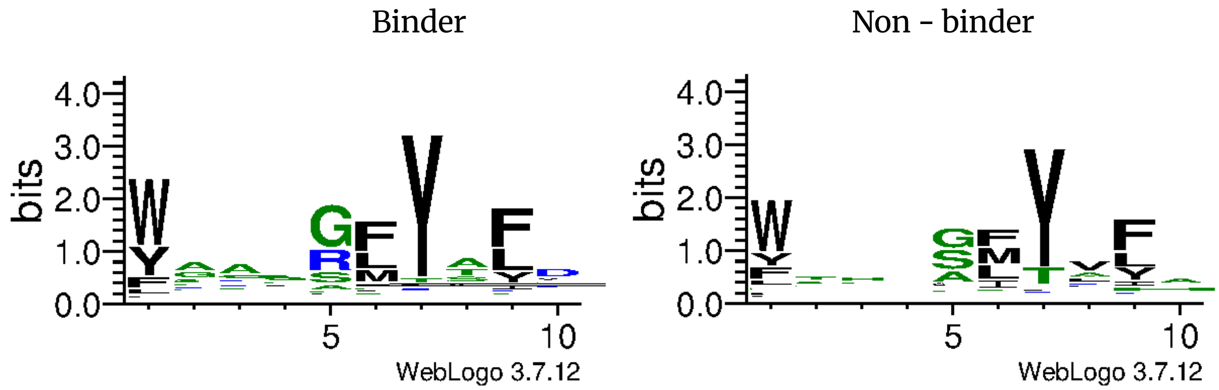


**Supplementary Figure 3: WebLogo residue distributions for HER2-large-aff CDRH3 sequences** - removed medium binders, binder (Left) and non-binder (Right). We can see 7th position is highly conserved.

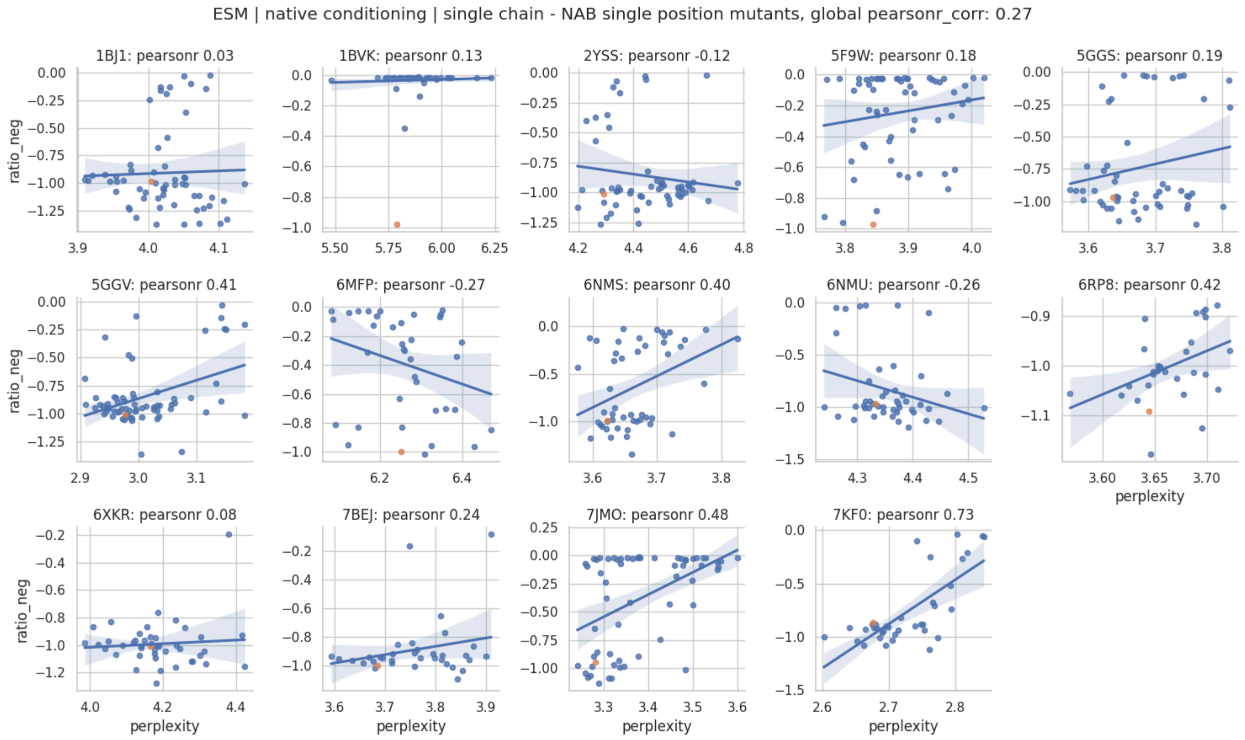


**Supplementary Figure 4. ESM-IF conditioned on native structure, no antigen.** Perplexity (x-axis) is plotted against the negative log ratio of the mutant/wt ELISA ratio (-1.0 is WT, closer to zero, worse binding).

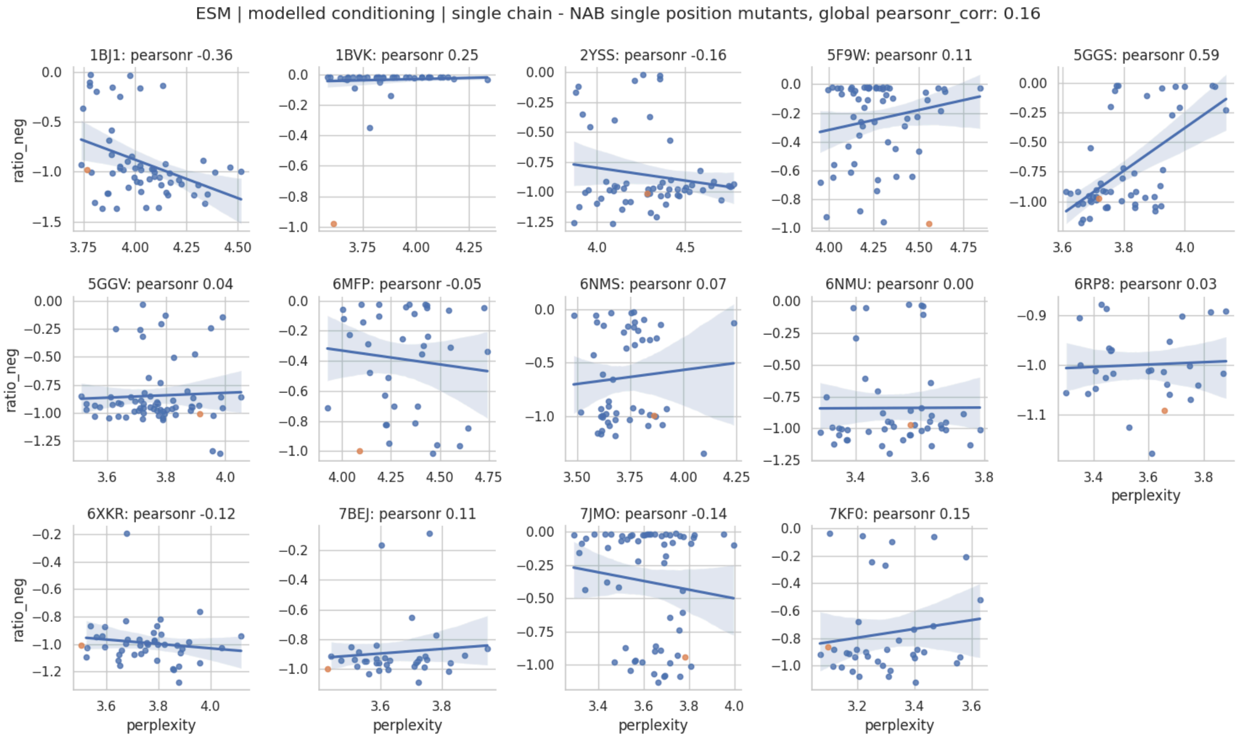


**Supplementary Figure 5. ESM-IF conditioned on model structure, no antigen.** Perplexity (x-axis) is plotted against the negative log ratio of the mutant/wt ELISA ratio (-1.0 is WT, closer to zero, worse binding).

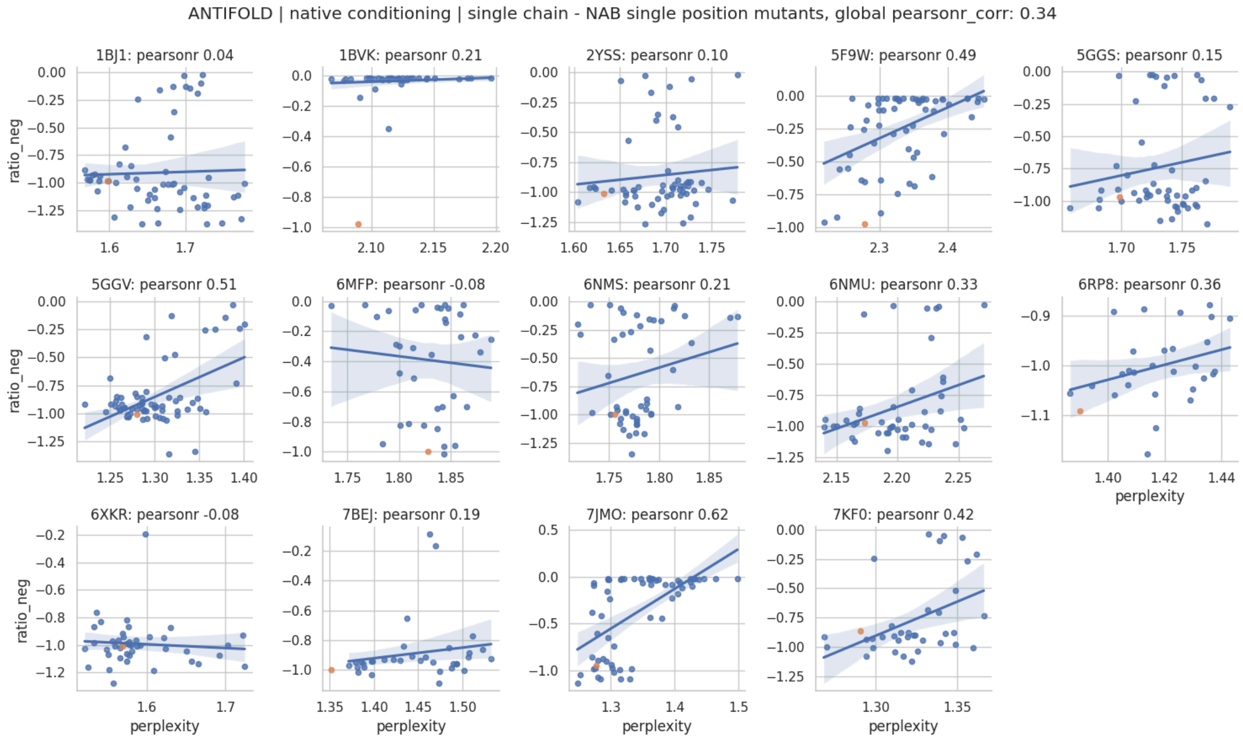


**Supplementary Figure 6. Antifold conditioned on native structure, no antigen.** Perplexity (x-axis) is plotted against the negative log ratio of the mutant/wt ELISA ratio (-1.0 is WT, closer to zero, worse binding).

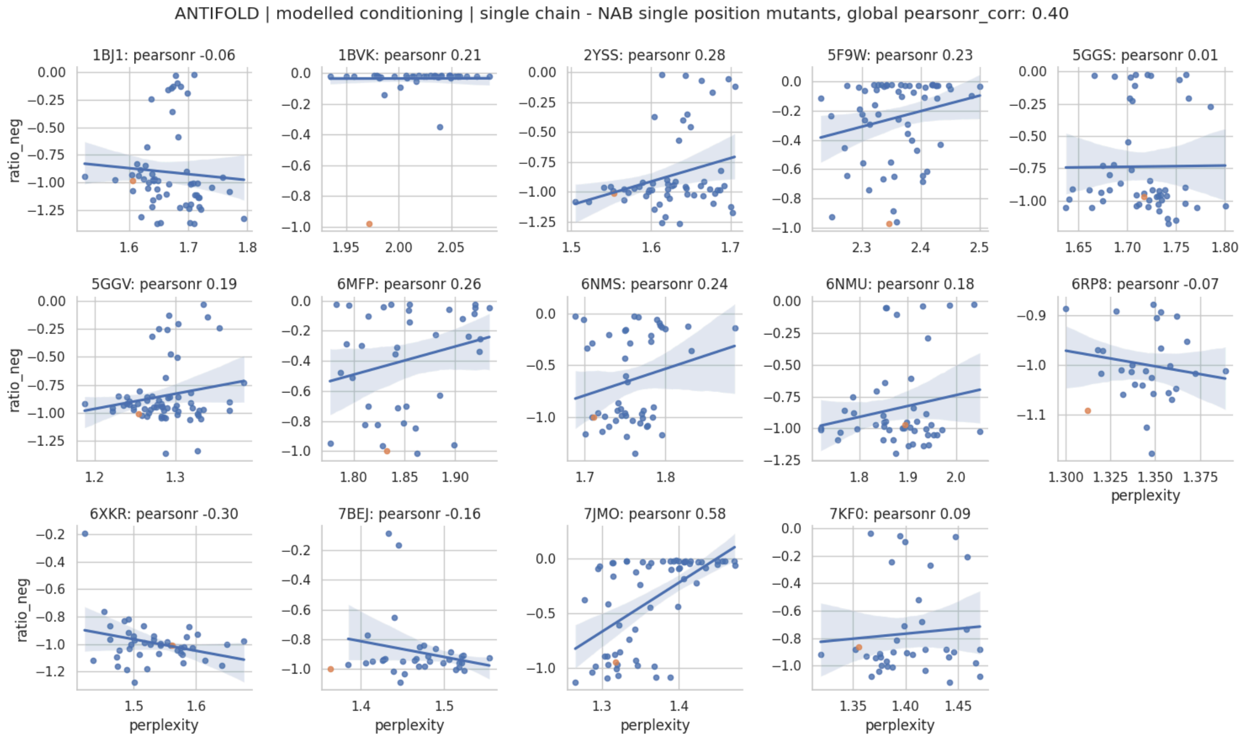


**Supplementary Figure 7. Antifold conditioned on model structure, no antigen.** Perplexity (x-axis) is plotted against the negative log ratio of the mutant/wt ELISA ratio (-1.0 is WT, closer to zero, worse binding).

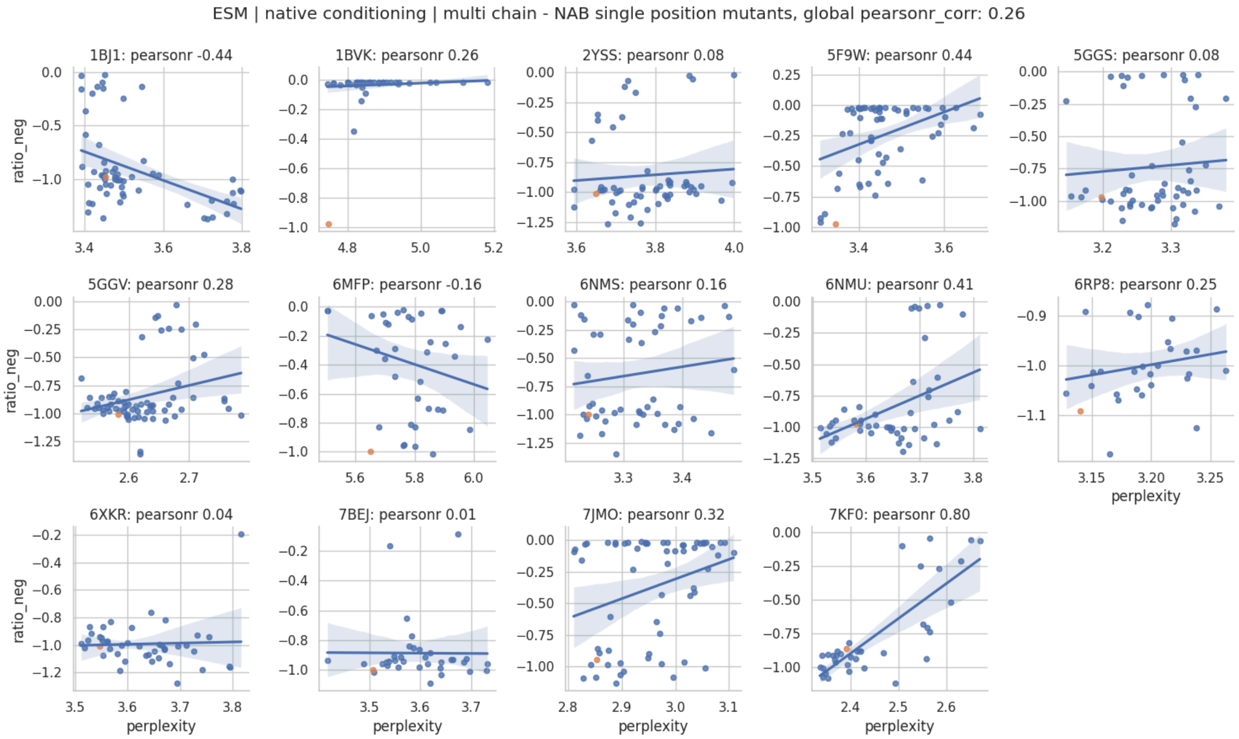


**Supplementary Figure 8. ESM-IF conditioned on native structure, with antigen.** Perplexity (x-axis) is plotted against the negative log ratio of the mutant/wt ELISA ratio (-1.0 is WT, closer to zero, worse binding).

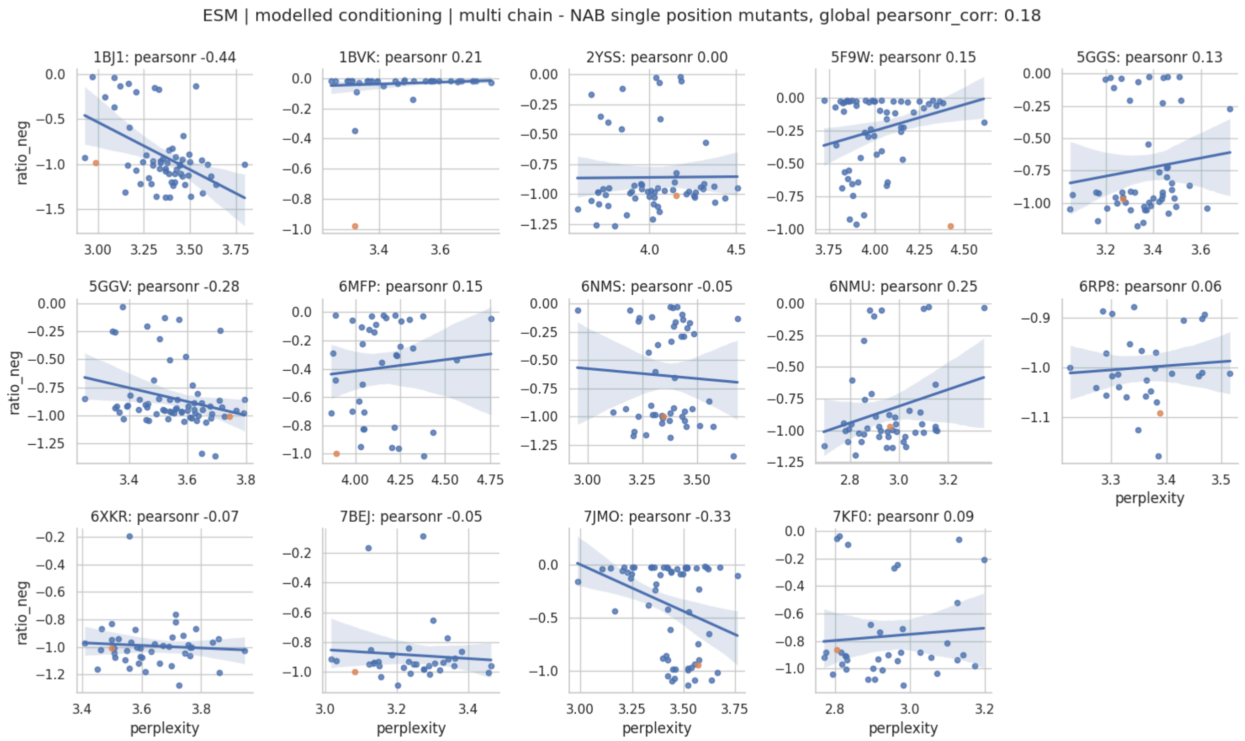


**Supplementary Figure 9. ESM-IF is conditioned on model structure, with antigen.** Perplexity (x-axis) is plotted against the negative log ratio of the mutant/wt ELISA ratio (-1.0 is WT, closer to zero, worse binding).

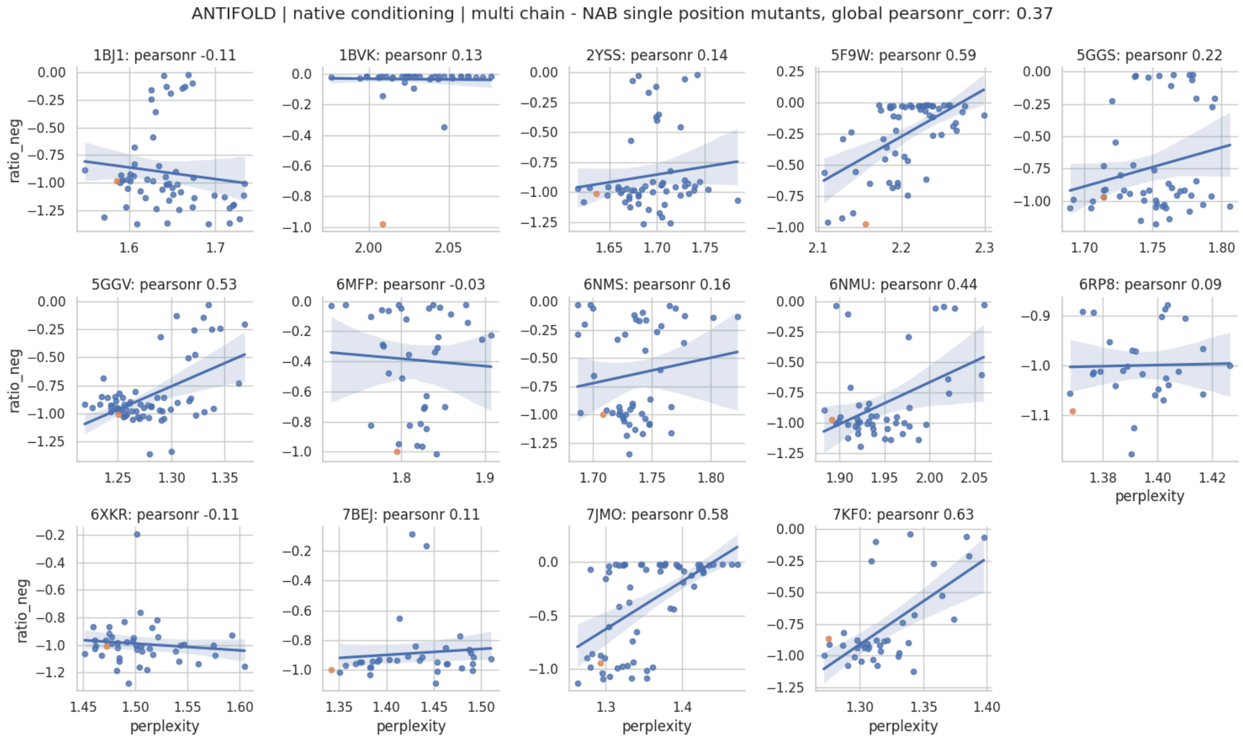


**Supplementary Figure 10. Antifold conditioned on native structure, with antigen.** Perplexity (x-axis) is plotted against the negative log ratio of the mutant/wt ELISA ratio (-1.0 is WT, closer to zero, worse binding).

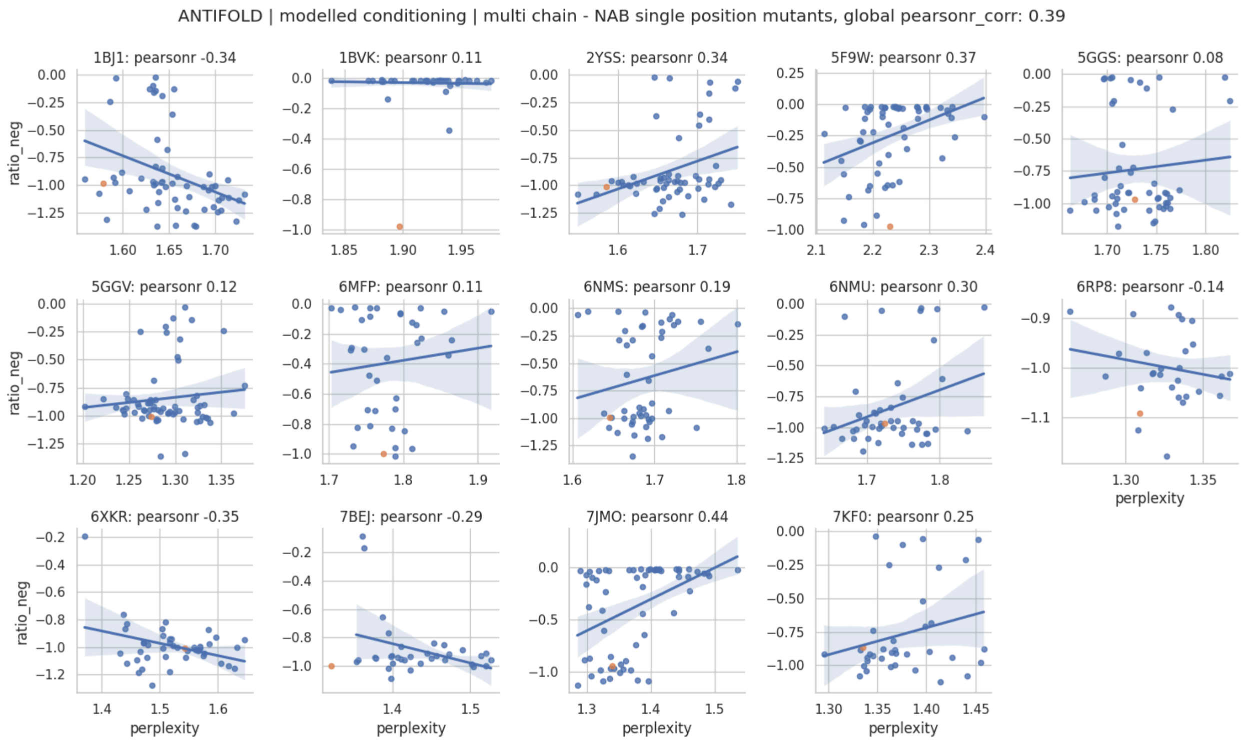


**Supplementary Figure 11. Antifold conditioned on model structure, with antigen.** Perplexity (x-axis) is plotted against the negative log ratio of the mutant/wt ELISA ratio (-1.0 is WT, closer to zero, worse binding).

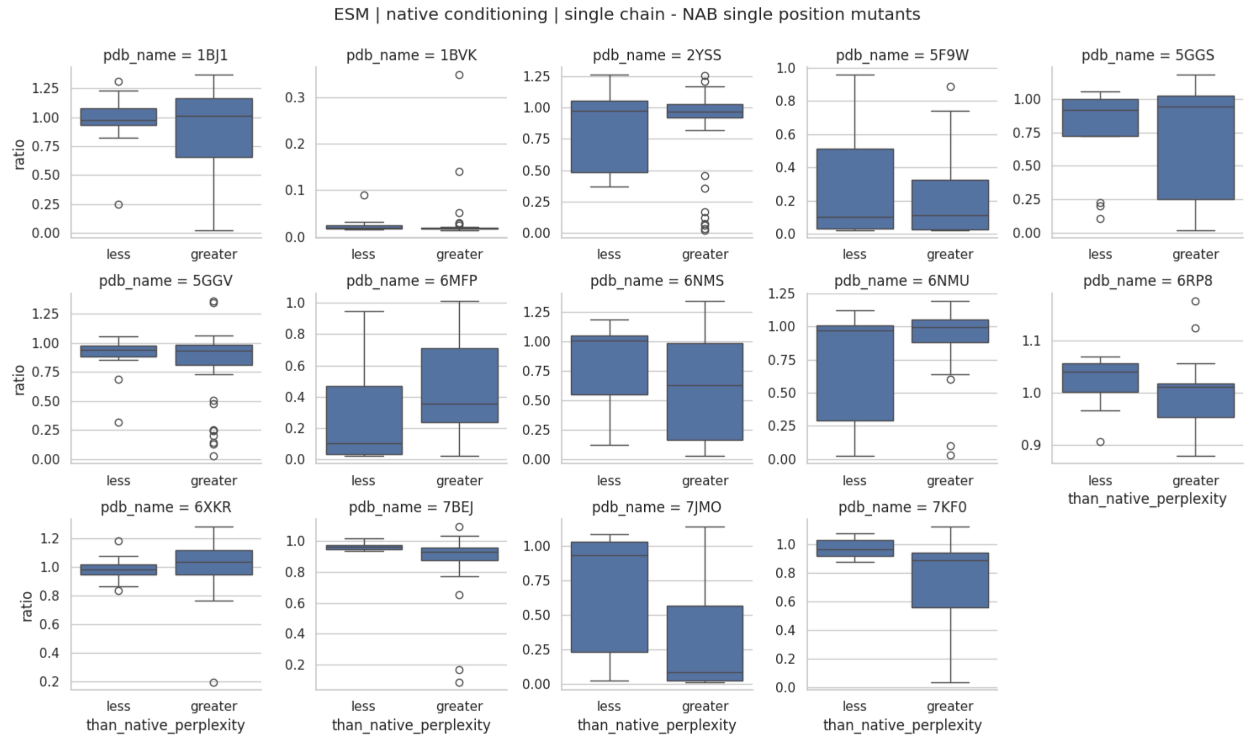


**Supplementary Figure 12. ESM-IF conditioned on native structure, no antigen.** 'Less' boxplots indicate perplexity scores smaller than Wild Type, whereas 'greater' indicate greater scores than wild type.

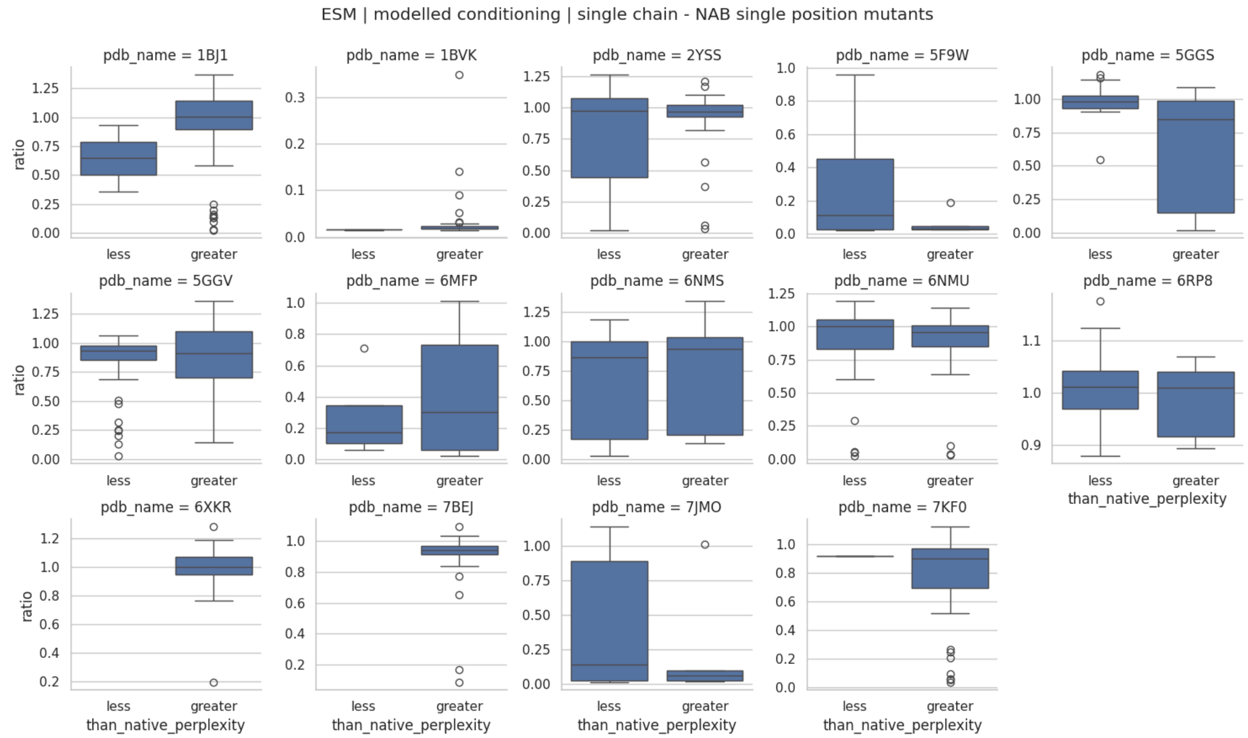


**Supplementary Figure 13. ESM-IF conditioned on model structure, no antigen.** 'Less' boxplots indicate perplexity scores smaller than Wild Type, whereas 'greater' indicate greater scores than wild type.

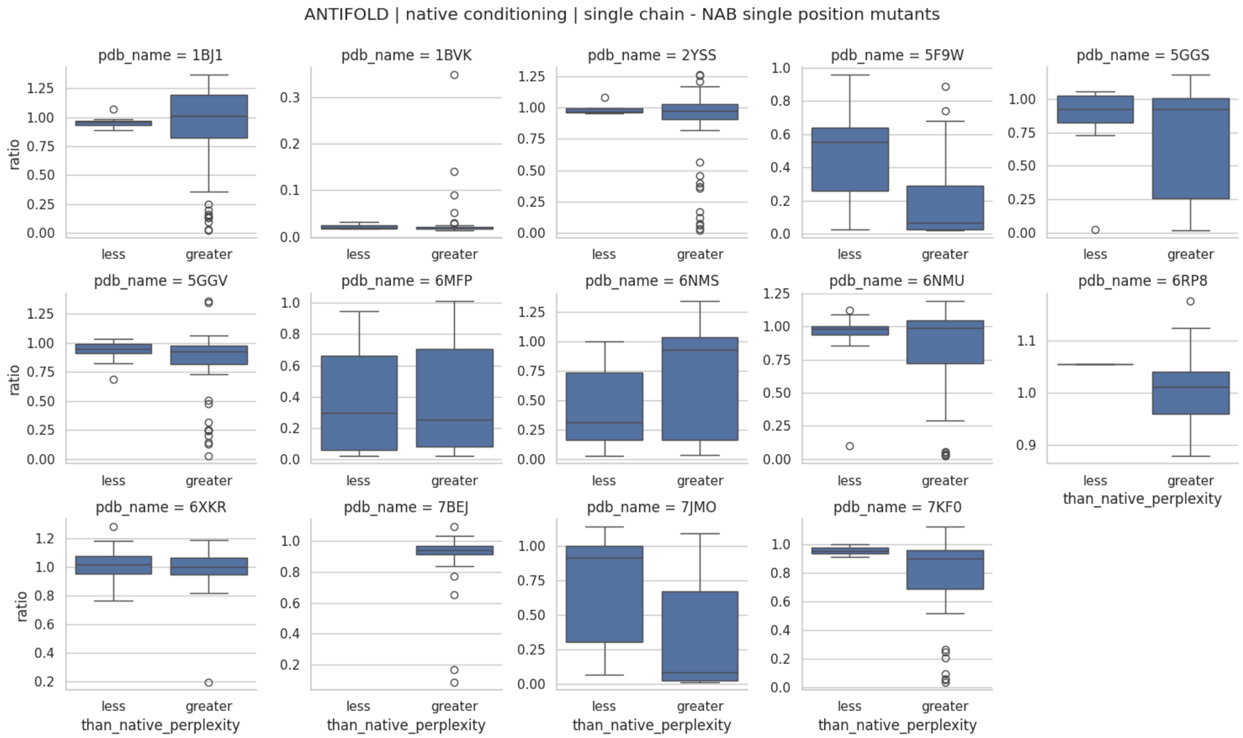


**Supplementary Figure 14. Antifold conditioned on native structure, no antigen.** 'Less' boxplots indicate perplexity scores smaller than Wild Type, whereas 'greater' indicate greater scores than wild type.

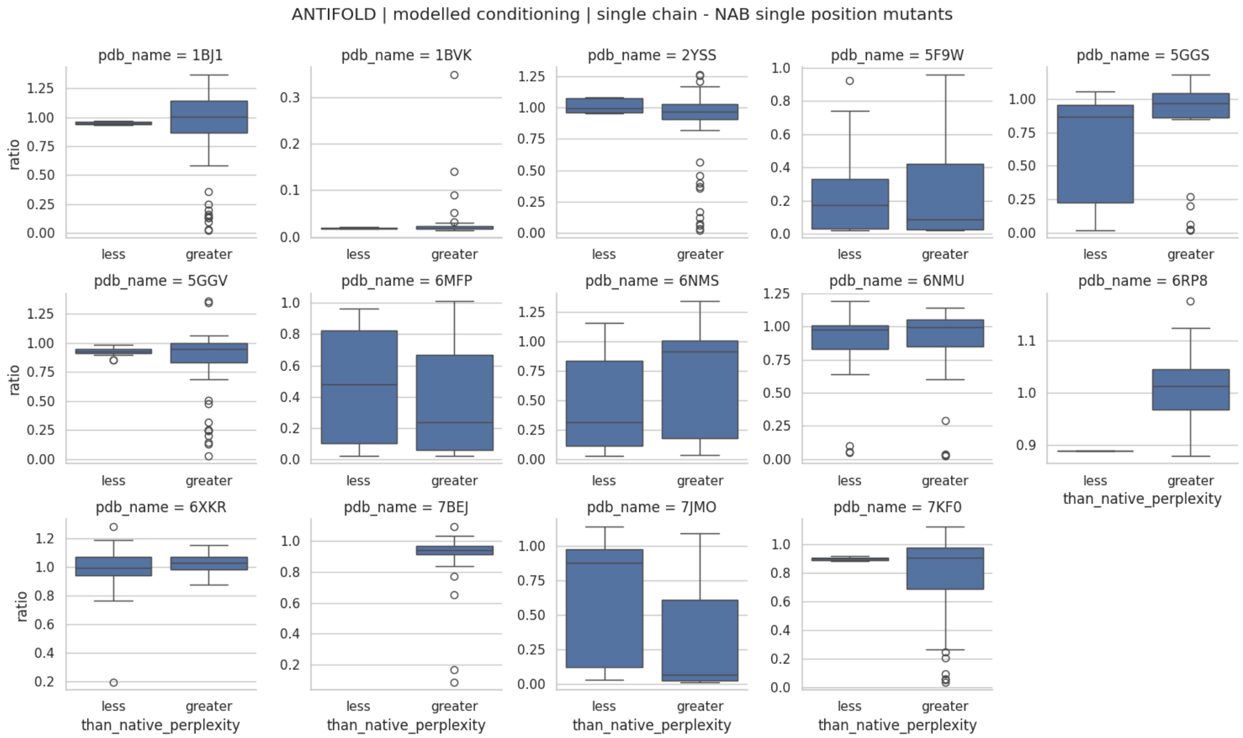


**Supplementary Figure 15. ESM-IF conditioned on model structure, no antigen.** 'Less' boxplots indicate perplexity scores smaller than Wild Type, whereas 'greater' indicate greater scores than wild type.

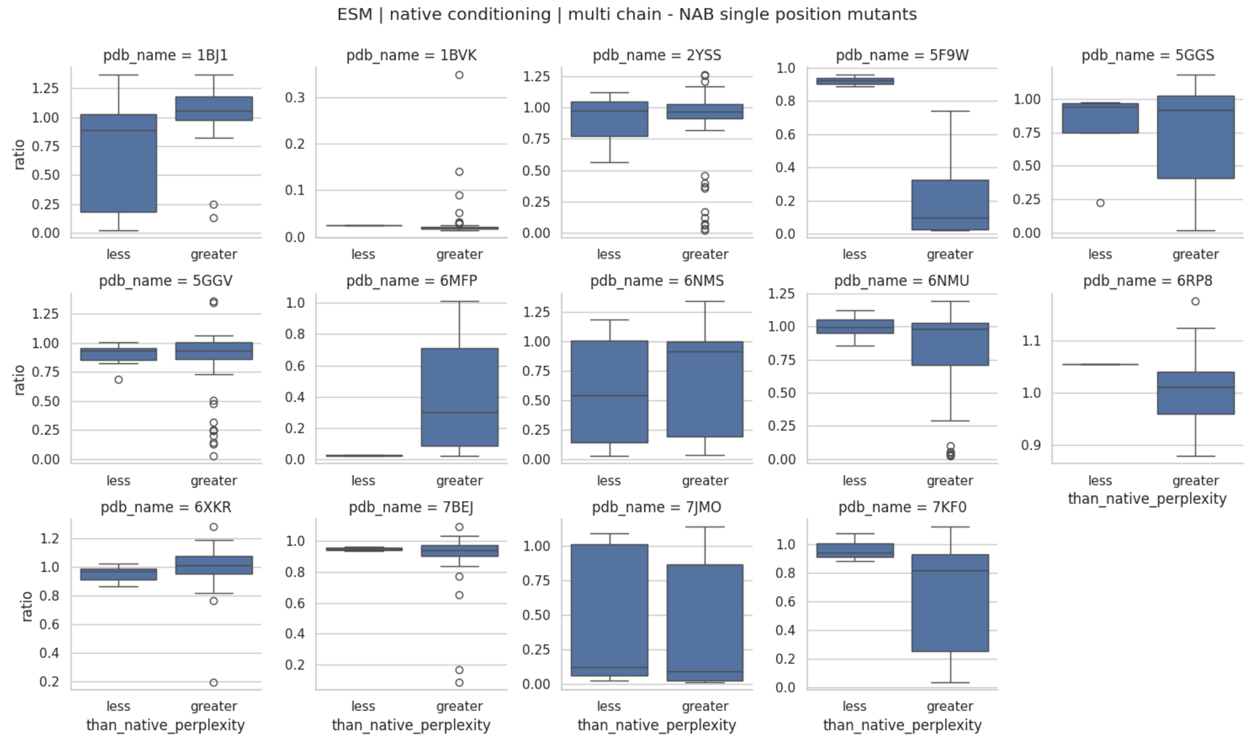


**Supplementary Figure 16. ESM-IF conditioned on native structure, with antigen.** 'Less' boxplots indicate perplexity scores smaller than Wild Type, whereas 'greater' indicate greater scores than wild type.

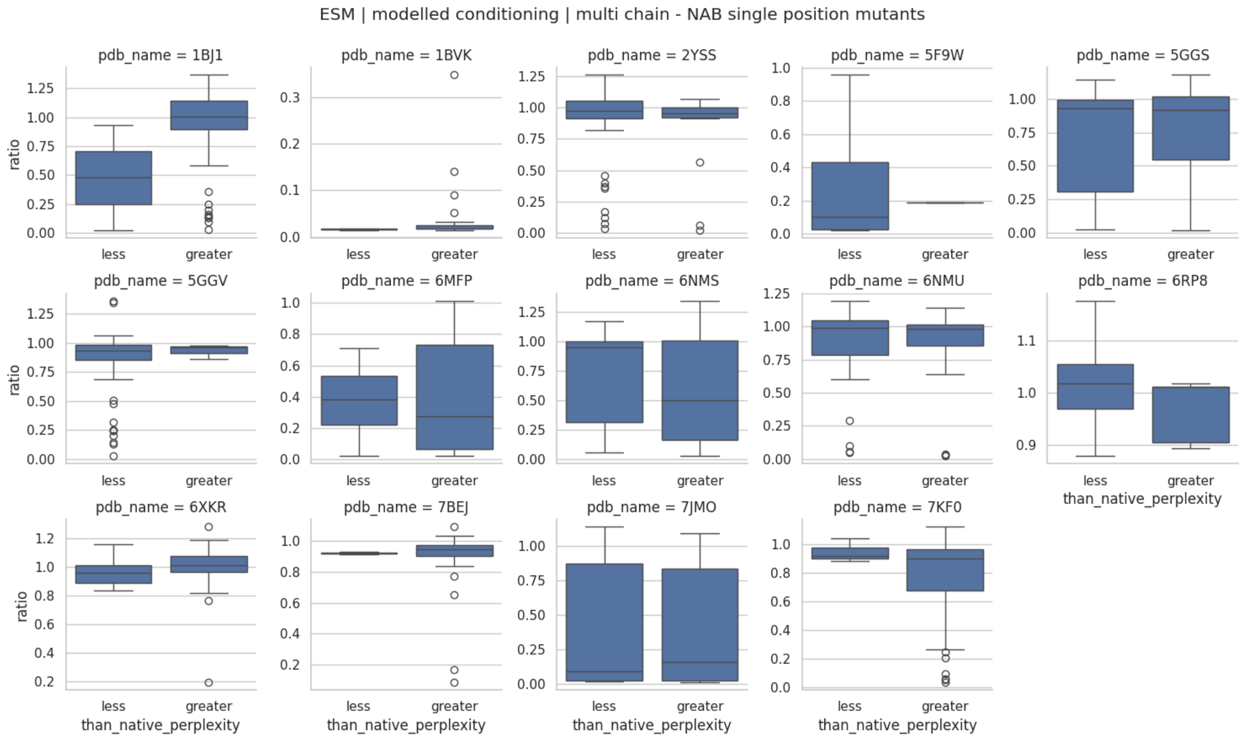


**Supplementary Figure 17. ESM-IF conditioned on model structure, with antigen.** 'Less' boxplots indicate perplexity scores smaller than Wild Type, whereas 'greater' indicate greater scores than wild type.

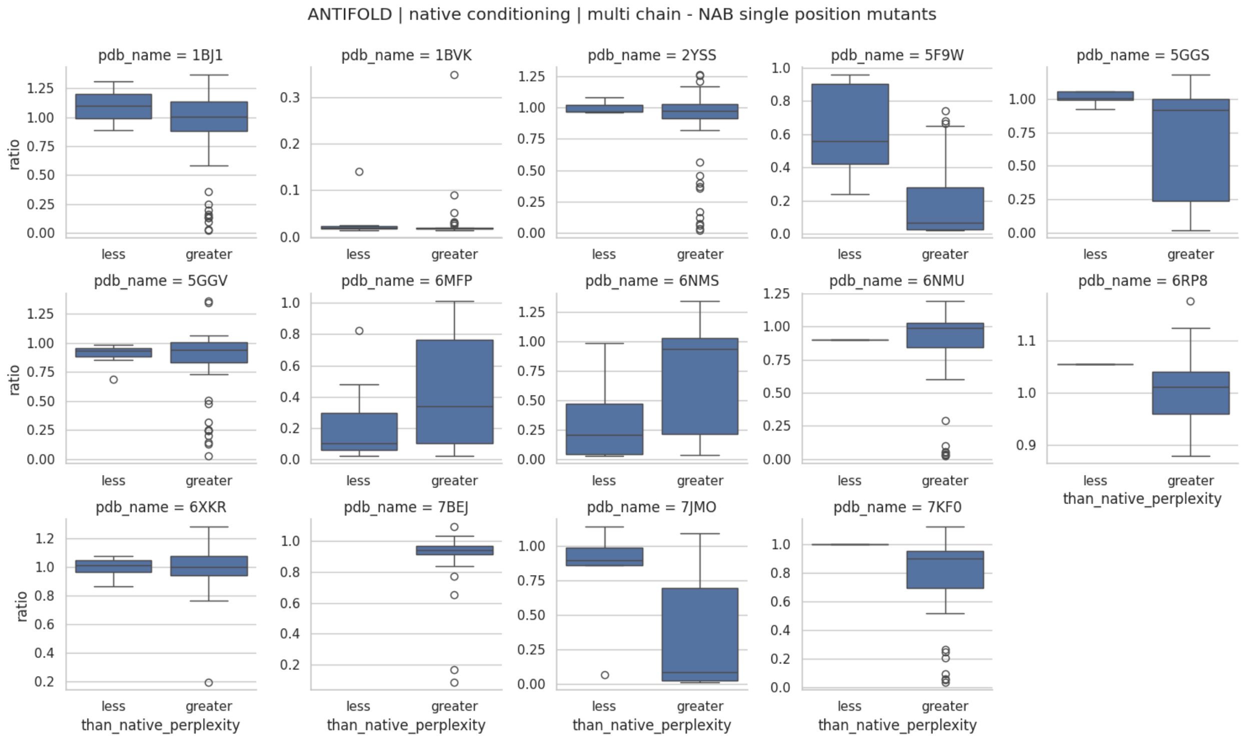


**Supplementary Figure 18. Antifold conditioned on native structure, with antigen.** ‘Less’ boxplots indicate perplexity scores smaller than Wild Type, whereas ‘greater’ indicate greater scores than wild type.

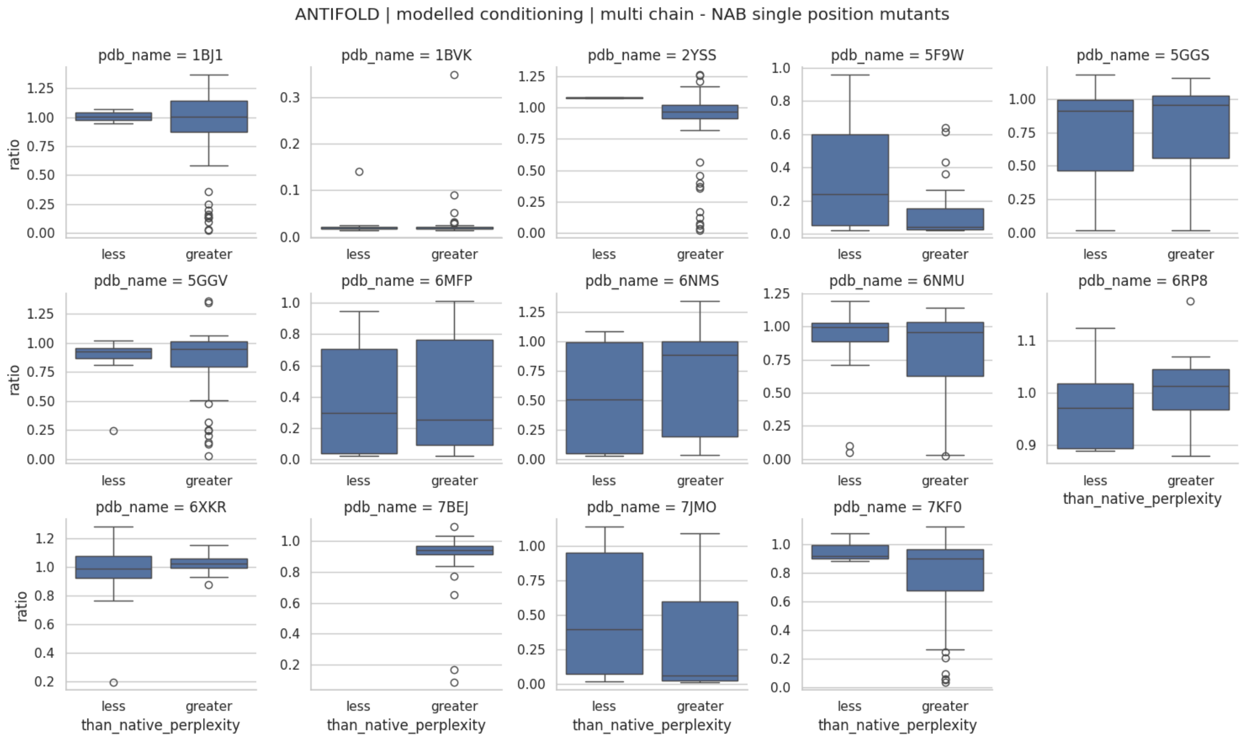


**Supplementary Figure 19. Antifold conditioned on model structure, with antigen.** 'Less' boxplots indicate perplexity scores smaller than Wild Type, whereas 'greater' indicate greater scores than wild type.

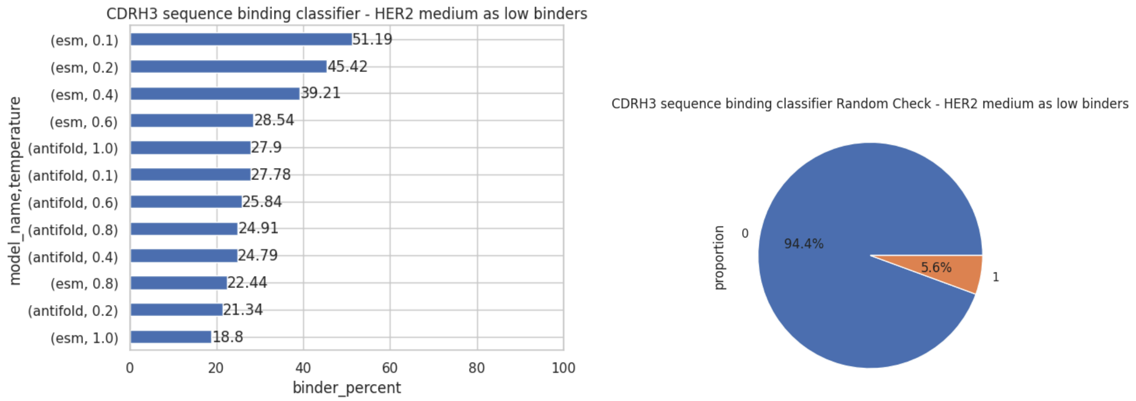


**Supplementary Figure 20: ESM-IF and Antifold sampled CDRH3 classification.** (Left) ESM-IF and Antifold sampled sequences classification by CNN model trained on CDRH3 sequences from HER2 dataset with medium labeled as low binders. Each bar corresponds to sequences generated using a given IF model on specific temperature setting. (Right) CDRH3 classifier sanity check - proportions of binder / non-binder classification on randomly generated CDH3 dataset.

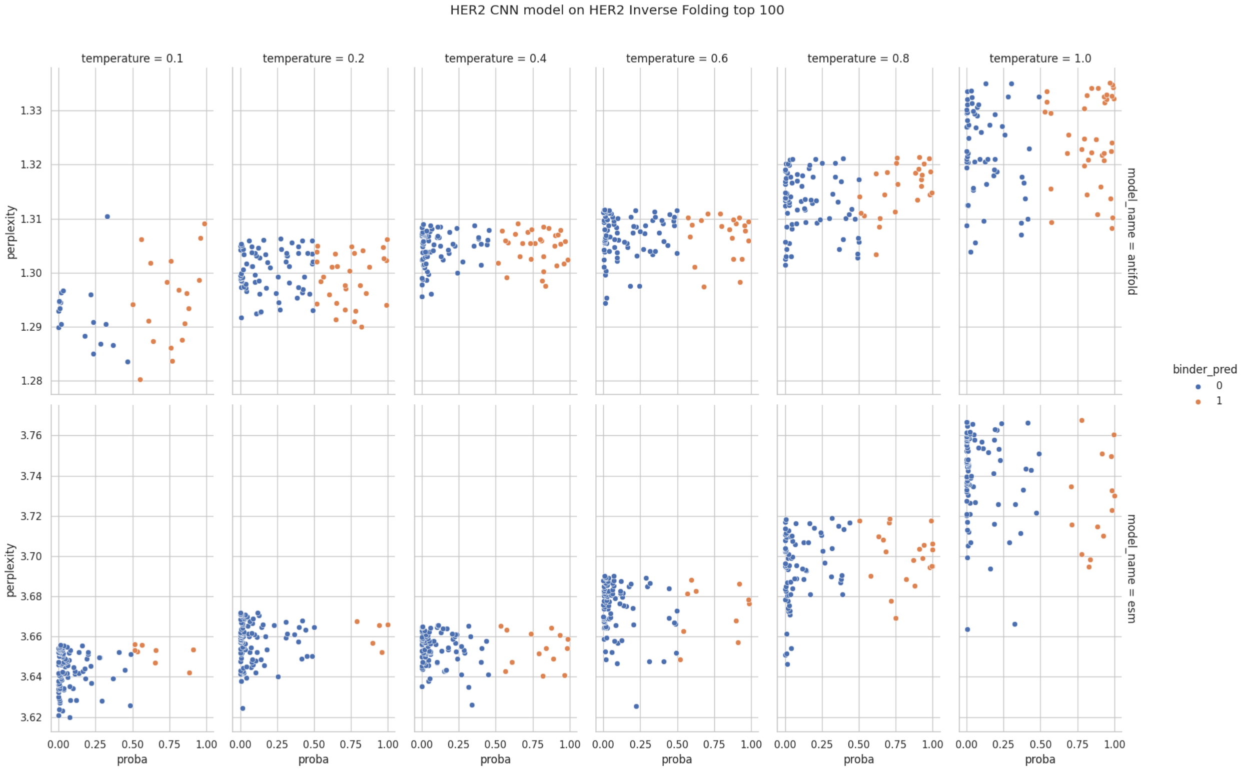


**Supplementary Figure 21: Perplexity distribution chart for ESM-IF and Antifold -** sampled sequences were scored using the same model they were generated with. Top 100 sequences for every model-temperature are displayed. Observations are colored by oracle classification CNN model trained on HER2 dataset with medium labeled as low binders.

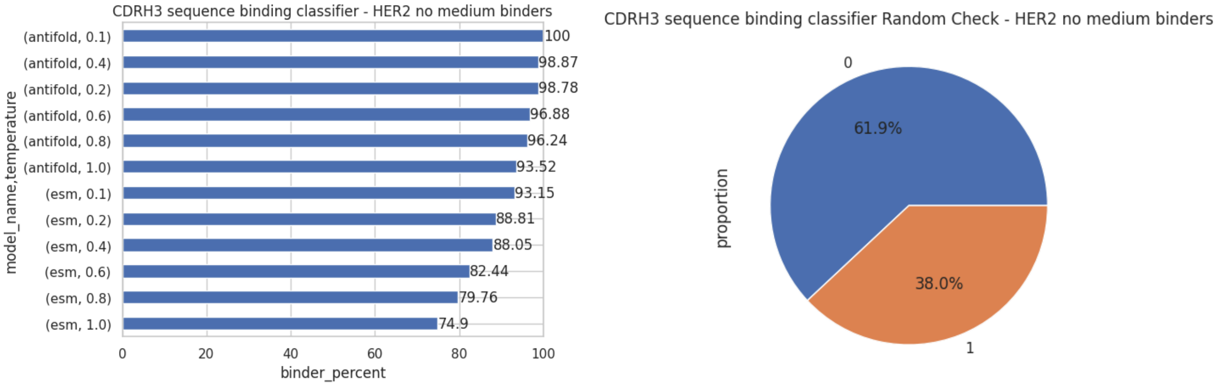


**Supplementary Figure 22: ESM-IF and Antifold sampled CDRH3 classification.** (Left) ESM-IF and Antifold sampled sequences classification by CNN model trained on CDRH3 sequences from HER2 dataset with removed medium binders. Each bar corresponds to sequences generated using a given IF model on specific temperature setting. (Right) CDRH3 classifier sanity check - proportions of binder / non-binder classification on randomly generated CDH3 dataset.

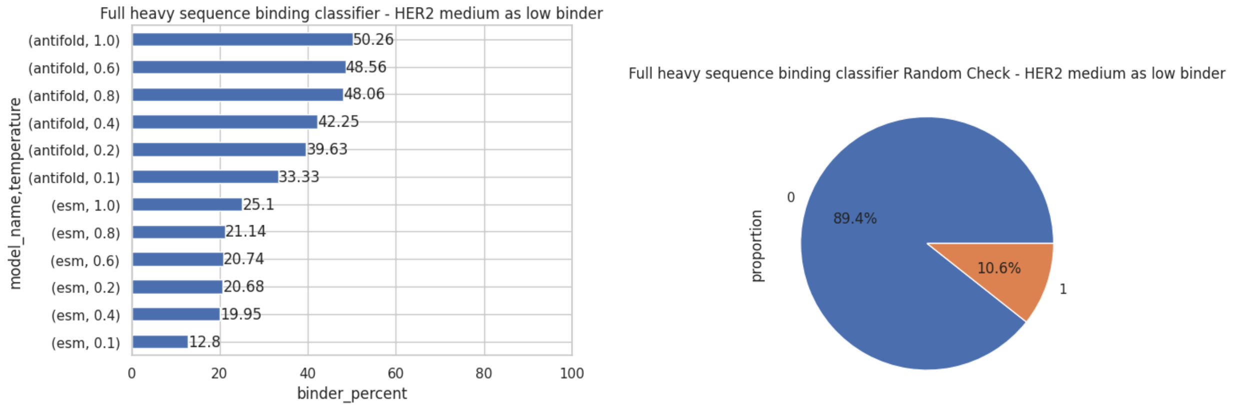


**Supplementary Figure 23: ESM-IF and Antifold sampled full sequence classification.** (Left) ESM-IF and Antifold sampled sequences classification by CNN model trained on full heavy HER2 dataset with medium labeled as low binders. Each bar corresponds to sequences generated using a given IF model on specific temperature settings. (Right) Full heavy sequence classifier sanity check - proportions of binder / non-binder classification on randomly generated CDH3 dataset.

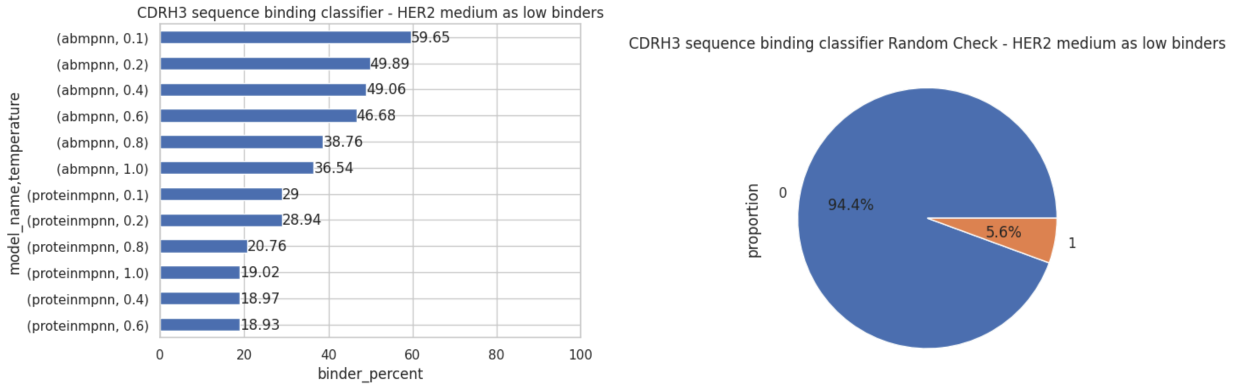


**Supplementary Figure 24: ProteinMPNN and AbMPNN sampled CDRH3 classification.** (Left) ProteinMPNN and AbMPNN sampled sequence classification by CNN model trained on CDRH3 sequences from HER2 dataset with medium labeled as low binders. Each bar corresponds to sequences generated using a given IF model on specific temperature setting. (Right) CDRH3 classifier sanity check - proportions of binder / non-binder classification on randomly generated CDH3 dataset.

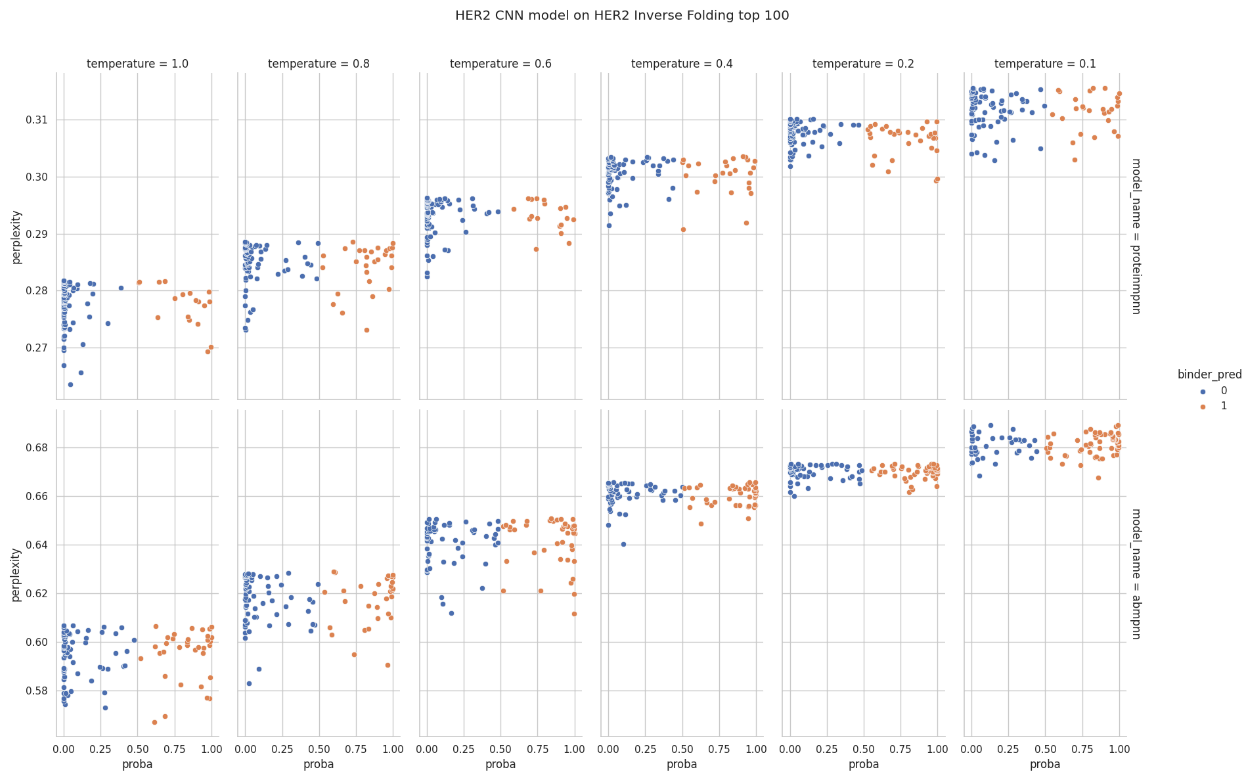


**Supplementary Figure 25: Perplexity distribution chart for ProteinMPNN and AbMPNN.** Sampled sequences were scored using the same model they were generated with. Top 100 sequences for every model-temperature are displayed. Observations are colored by oracle classification CNN model trained on HER2 dataset with medium labeled as low binders.

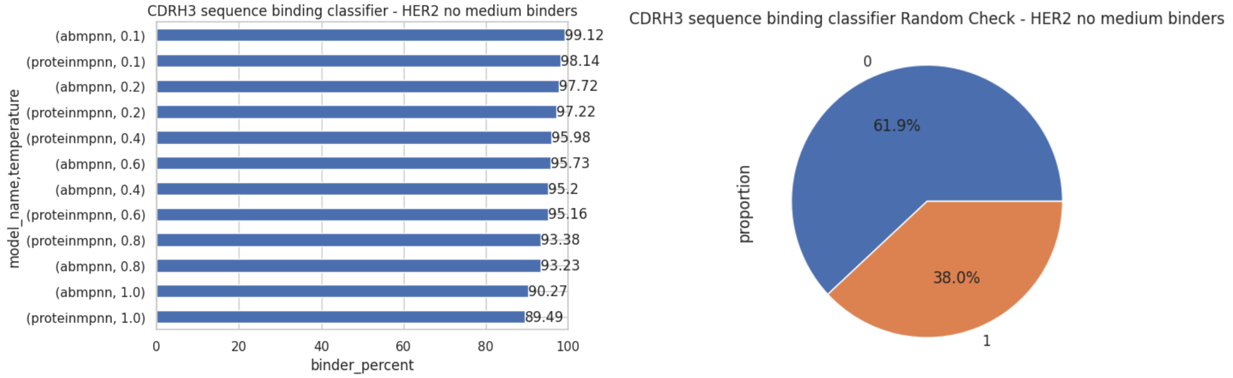


**Supplementary Figure 26: ProteinMPNN and AbMPNN sampled CDRH3 classification.** (Left) ProteinMPNN and AbMPNN sampled sequences classification by CNN model trained on CDRH3 sequences from HER2 dataset with removed medium binders. Each bar corresponds to sequences generated using a given IF model on specific temperature setting. (Right) CDRH3 classifier sanity check - proportions of binder / non-binder classification on randomly generated CDH3 dataset.

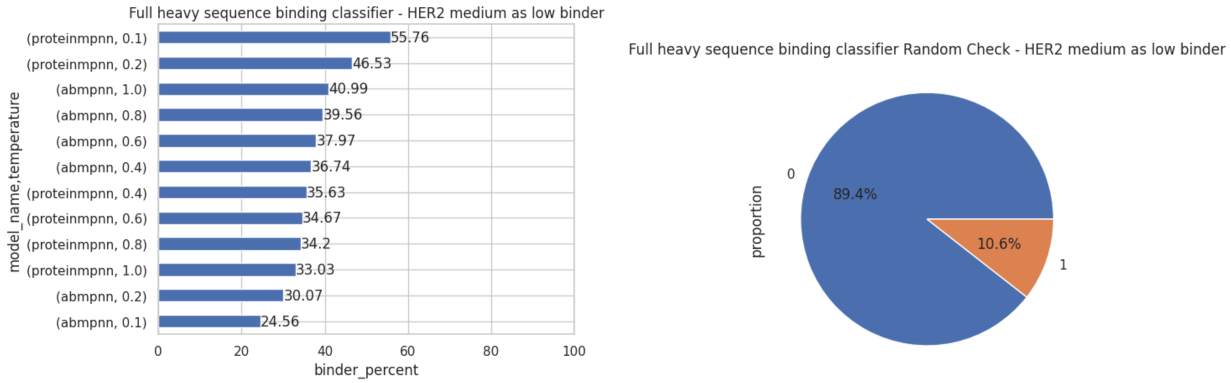


**Supplementary Figure 27: ProteinMPNN and AbMPNN sampled full sequence classification.** (Left) ProteinMPNN and AbMPNN sampled sequences classification by CNN model trained on full heavy HER2 dataset with medium labeled as low binders. Each bar corresponds to sequences generated using a given IF model on specific temperature setting. (Right) Full heavy sequence classifier sanity check - proportions of binder / non-binder classification on randomly generated CDH3 dataset.
